# Galectin and Myc enable cochlear progenitor expansion in vitro and in vivo

**DOI:** 10.64898/2026.06.03.729765

**Authors:** Marie Kubota, Julia M. Abitbol, Paul K. Lee, Maggie S. Matern, Sonia Bustos Barocio, Taha A. Jan, Alan G. Cheng, Stefan Heller

## Abstract

The neonatal cochlear epithelium harbors regenerative capacity, attributable to the transient greater epithelial ridge (GER). After injury, GER cells re-enter the cell cycle, migrate into the damaged organ of Corti, and differentiate into sensory or supporting cells *in vivo*; in culture, they proliferate to form inner ear organoids. The mechanisms underlying this competence remain unclear. Here, we generated organoids from mouse GER cells and performed single-cell transcriptomics at organoid initiation. Analysis revealed extracellular matrix reorganization with prominent involvement of galectins. Pharmacological inhibition reduced organoid growth, whereas overexpression of galectin-1 or Myc enhanced growth, identifying both factors as regulators of cochlear progenitor expansion. Moreover, Myc overexpression conferred organoid-forming capacity in post-neonatal cochlear epithelial cells after hearing onset, thereby extending the time window for regenerative competence. *In vivo*, organ of Corti ablation induced galectin-1 upregulation and GER proliferation, suppressed by galectin-1 blockade, establishing a model to rekindle proliferative potential in mature cochlear cells.

## Introduction

Loss of cochlear hair cells is a major cause of hearing loss in mammals, resulting from aging, ototoxic drugs, genetic predispositions, or noise exposure. In the mature cochlea, non-sensory cells such as organ of Corti supporting cells and adjacent epithelial cells are mitotically quiescent and lack regenerative capacity after hair cell demise. However, ectopic activation of specific transcription factors and signaling pathways can induce regenerative responses in these cells ^1–4^. This strategy is particularly effective in neonatal mice, where cochlear non-sensory cells can proliferate and generate hair cells both *in vitro* and *in vivo* ^1,5,6^. A particularly plastic progenitor cell population for the organ of Corti is the adjacent greater epithelial ridge (GER) ^7–10^. The GER is a transient neonatal cell population that disappears during the first postnatal week before the onset of hearing ^11^. We previously showed that fluorescence-activated cell sorting (FACS)-enriched GER cells display robust regenerative capacity in culture. This cell population represented the most potent cochlear progenitors, capable of proliferating and forming otic organoids that develop sensory epithelia containing nascent hair cells and supporting cells ^9^.

Given the robust progenitor activity of GER cells, we hypothesized that transcriptional changes accompanying the transition into organoid growth could identify regulators of proliferative competence in cochlear epithelial cells. We therefore performed single-cell RNA sequencing of GER-derived organoids during the early stages of organoid formation and identified candidate pathways associated with this transition. We first tested candidate pathways pharmacologically for effects on organoid formation and growth and found that inhibition of galectin-1, galectin-3, or Myc suppresses organoid growth.

Using adeno-associated virus (AAV)-mediated overexpression, we then demonstrated that galectin-1 and Myc enhance organoid growth from GER cells. When overexpressed in cochlear epithelial cells isolated from mice older than two weeks, which lack GER, Myc increased proliferation, providing proof of concept that post-neonatal cochlear cells can be induced to proliferate in the organoid system using genes that are functionally linked to GER cells’ regenerative abilities. *In viv*o, we detected increased galectin-1 and Myc expression in the GER using an Lgr5-diphtheria toxin receptor (DTR)-based mouse cochlear cell ablation model, in which GER cells enter the cell cycle following the ablation of Lgr5-positive supporting and epithelial cells ^12^. Inhibition of galectin-1 decreased GER cell proliferation after cell ablation, revealing that galectin signaling is required for damage-induced GER cell proliferation *in vivo*. Our study provides a transcriptomic atlas of GER cell proliferation in organoids and demonstrates that selected pathways contribute functionally to organoid growth. Moreover, we show that these pathways are actively engaged in the regenerative response following *in vivo* injury. We propose that GER-derived organoid systems provide a platform for developing strategies to regenerate mature cochlear cells.

## Results

### Single cell collection from purified GER cell-derived organoids

GER cells robustly give rise to otic organoids that develop epithelial structures harboring nascent hair cells and supporting cells within three weeks ^9^. To capture dynamic changes in gene expression during organoid growth, we performed single-cell RNA sequencing of GER-derived organoids (GER-organoids). We analyzed organoids that formed after one day, three days, and seven days *in vitro*, hypothesizing that these time points encompass the initial phase of GER cell proliferation, and thus allow identification of genes involved in this process.

GER cells were purified with FACS from postnatal day (P) 2 mice (*Sox2^GFP/+^; Fgfr3^CreERT2/+^; Ai14^tdTomato/+^*) by sorting a cell population expressing high levels of GFP and lacking tdTomato fluorescence (**Figure 1A**) ^9,13,14^. We have previously developed a method for collecting GER cells by the intensities of two fluorescent reporters expressed in the mouse line with FACS (Gate 1: GFP-high/ tdTomato-negative; Gate 2: GFP-low/ tdTomato-negative; Gate 3: GFP-high/ tdTomato-high; Gate 4: GFP-low/ tdTomato-high). The cells from Gate 1 to Gate 4 were sorted into designated wells of 96-well plates, and single-cell RNA sequencing was performed on these cells (**Figure 1B**). Each cell’s well position was retained during analysis, allowing assignment to its corresponding FACS gate. Clustering analysis identified the cell subtypes present in each FACS gate. As a result, we confirmed that GER cells are highly enriched in Gate 1 with a purity exceeding 90% ^9,13^ (**Figure 1B**). The FACS-purified GER cells from Gate 1 were seeded and cultured for seven days to form organoids.

**Figure 1.**
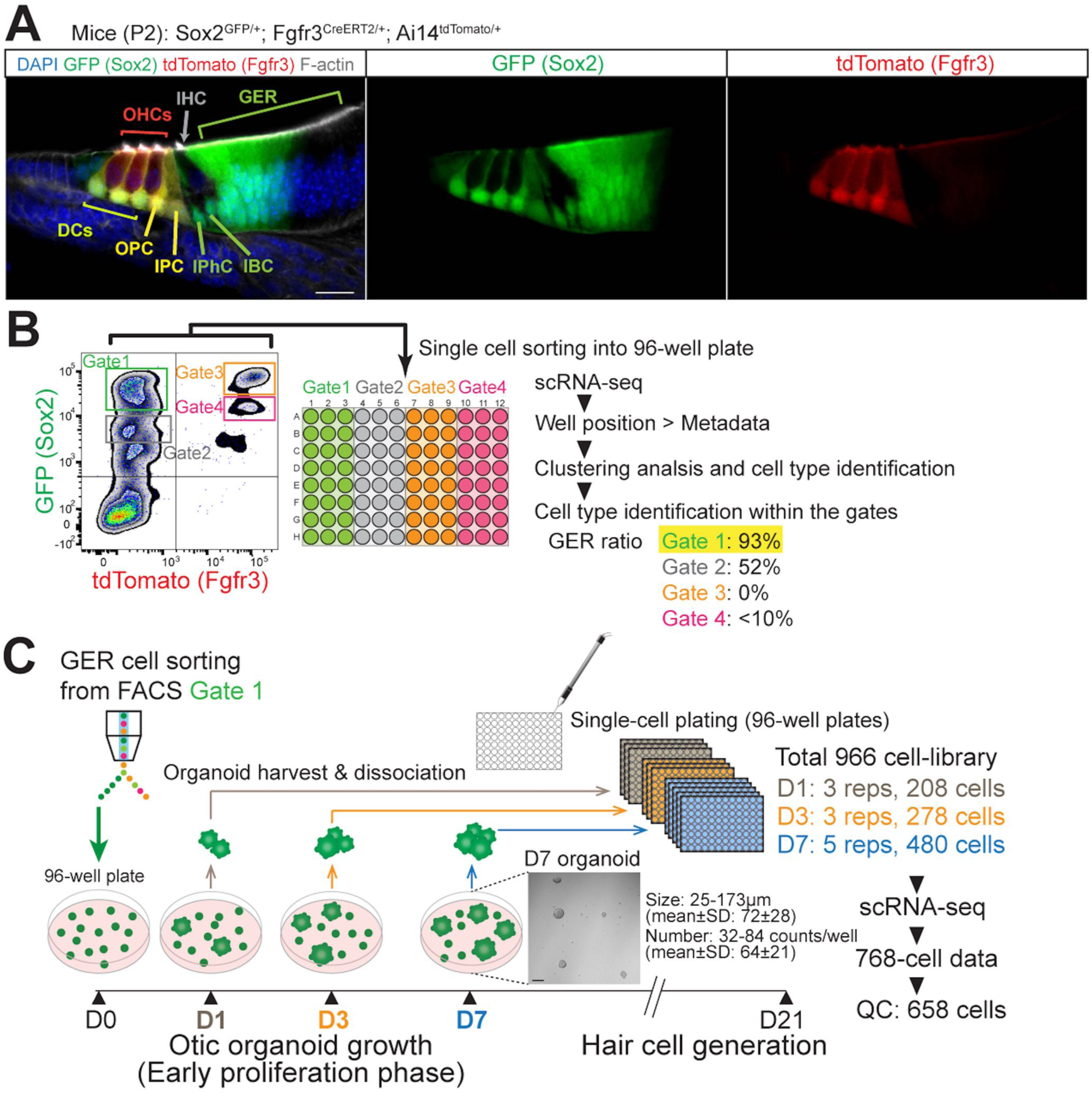
Single-cell RNA sequencing of GER cell-derived organoids. **(A)** Fluorescent reporter gene expression in a P2 organ of Corti vibratome section of an Fgfr3-Cre-ERT2/Ai14-tdTomato/Sox2-GFP transgenic mouse. Scale bar: 20 µm. DCs: Deiters’ cells; OPC: outer pillar cell; IPC: inner pillar cell; IPhC: inner phalangeal cell; IBC: inner border cell; IHC: inner hair cell; OHCs: outer hair cells; GER: greater epithelial ridge. **(B)** Enrichment strategy of GER cells from Fgfr3-Cre-ERT2/Ai14-tdTomato/Sox2-GFP transgenic mouse. Dissociated P2 cochlear floor cells were sorted from individual FACS gates (Gates 1-4) into defined positions in 96-well plates and subjected to single-cell RNA sequencing (scRNA-seq). Well-position metadata were then matched with cell type identities defined by clustering analysis to quantify the proportion of GER cells in each gate. A cell population expressing high GFP and lacking tdTomato fluorescence (Gate 1) contained GER cells with 93% purity ^9,13^. **(C)** Single-cell harvesting and sequencing from GER-derived organoids. FACS-enriched GER cells from Gate 1 were seeded at 2,000 cells per well for organoid growth in suspension culture for 7 days. Organoids were harvested on culture days 1 (D1), 3 (D3), and 7 (D7). A representative image of D7 organoids is shown (scale bar: 200 µm). Manual harvesting of single cells from dissociated organoids using a glass pipette of 50-60 µm diameter yielded a total library of 966 cells. Of these, scRNA-seq data were obtained from 768 cells, and quality control filtering retained 658 cells for downstream analysis.

At culture day seven, from 2,000 FACS-enriched GER cells seeded per well, we counted 64 ± 21 (mean ± SD) organoids per well with an average diameter of 72 ± 28 µm (mean ±SD) (**Figure 1C**). Organoid numbers and sizes were both smaller at earlier timepoints, resulting in insufficient cell concentrations for single-cell library preparation when we used bulk cell suspension methods. FACS-based sorting was also not feasible at these low cell numbers.

To overcome this challenge, dissociated organoid cells on culture days one, three, and seven were manually collected in 96-well plates using 50-60 µm diameter glass micropipettes and processed for single-cell RNA sequencing using the Smart-seq2 platform ^15^. Out of 966 cell-libraries (208, 278, and 480 cells from day one, three, and seven, respectively), single-cell RNA sequencing data were obtained for 768 cells, with 658 cells passing quality control and used for downstream analysis (**Figure 1C**).

### Identification of candidate genes that mediate otic organoid growth

The 658 cells were subjected to data integration, normalization, and clustering with Seurat ^16^. We identified seven distinct cell clusters in the dataset (**Figure 2A, Table S1**). Cells obtained from day one organoids were predominantly localized in cluster 6. Clusters 4 and 5 contained cells from both day one and day three organoids. Cells from day three and day seven organoids co-localized within clusters 1 and 3, while cluster 2 primarily comprised cells obtained from day seven organoids (**Figure 2A, B**). The expression of the proliferation marker Kiel 67 (*Mki67*) identified proliferating cells predominantly in clusters 1 and 3 (**Figure 2C**). Consistent with *Mki67* upregulation, Ingenuity Pathway Analysis (IPA) ^17^ revealed that genes associated with cell division were upregulated in clusters 1 and 3, which mainly contained cells from days three and seven. Furthermore, *Mki67* mRNA expression was not detected in cluster 2, which consisted of day seven cells. These dynamics suggest that organoid cells enter a proliferative state by day three, whereas a subgroup of day seven cells (cluster 2) did not proliferate. IPA identified pathways such as nerve growth factor signaling and Notch signaling in the non-proliferative cells (**Table S2)**. Because proliferation emerged between days one and three, we reasoned that transcriptional states present during this interval could reveal pathways associated with the initiation of organoid growth. We therefore focused on clusters 4, 5, and 6. Cluster 6 contained a major population of D1 cells and was transcriptionally distinct from cluster 7, which also consisted predominantly of D1 cells. Based on these transcriptional differences, we interpreted cluster 7 as representing an earlier baseline D1 state, whereas cluster 6 represented cells beginning to transition toward the subsequent organoid growth and proliferative states. Clusters 4 and 5 contained cells from both D1 and D3 and were likewise positioned within this onset-associated transition. We therefore used clusters 4-6 for subsequent differential-expression analysis.

**Figure 2.**
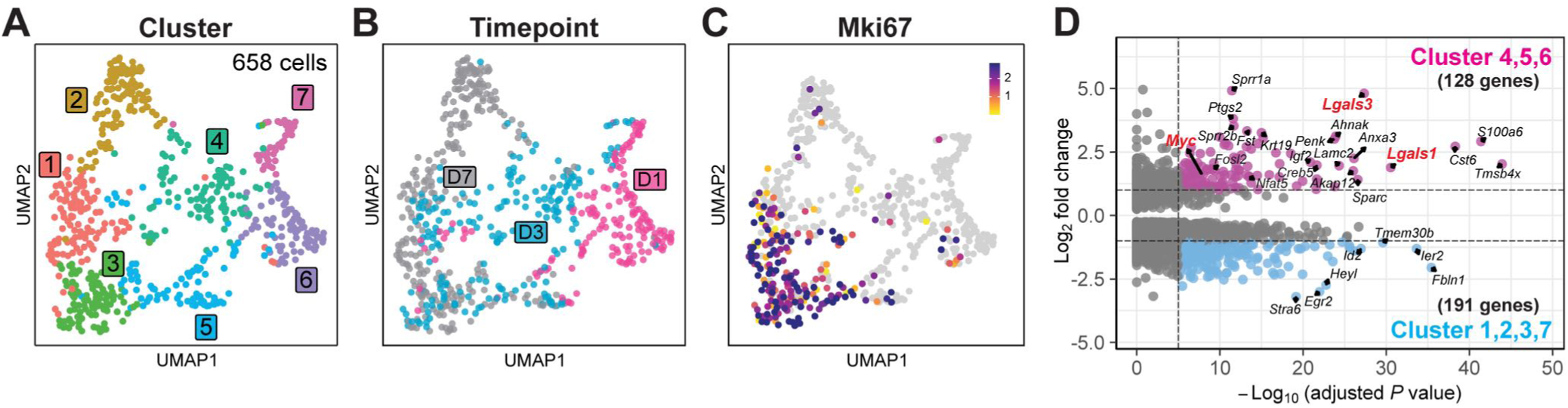
Single-cell RNA sequencing of GER-derived organoids reveals proliferating cell clusters. **(A)** Seurat data analysis of 658 cells, visualized by Uniform Manifold Approximation and Projection (UMAP), identified seven distinct clusters. Each dot represents a cell, colored by cluster. **(B)** UMAP projection, visualized by timepoint, showing the origin of cells from D1, D3, and D7 organoids. **(C)** Robust expression of mRNA encoding proliferation marker Kiel 67 (*Mki67*) was detected in the cells represented by clusters 1 and 3. **(D)** Volcano plot showing differentially expressed genes between the cells at the onset of proliferation (clusters 4, 5, and 6) and the remaining clusters (1, 2, 3, and 7). Cluster selection was based on transcriptional state rather than collection time points alone. Thresholds are indicated with dotted lines and were set to an absolute average log2 fold change (FC) > |1| and adjusted p-value < 1E-5, corresponding to -log_10_ (adjusted P) > 5. The Wilcoxon Rank Sum test was used for the comparison.

To identify putative cell growth-promoting effectors, we evaluated differentially expressed genes in these three clusters. The list of upregulated genes relative to the other clusters (clusters 1, 2, 3, and 7) (**Figure 2D, Table S3**) indicated active production of extracellular matrix (ECM) proteins, which aligned with IPA results showing the activation of signaling related to collagen biosynthesis and ECM-cell surface interactions on day one (cluster 6), followed by cell junction organization from day one to three (cluster 4) (**Table S2**).

An independent clustering approach based on hierarchical spectral clustering identified 13 clusters (**Figure S1A, Table S4**). Using similarity weighted non-negative embedding (SWNE) ^18^, which considers latent factors in two-dimensional projections, the organoid cells were arranged progressively from day one to day three and day seven. Proliferating cells, indicated by the upregulation of *Mki67* gene expression, were found in six clusters, primarily consisting of day three cells and a subset of day seven cells (**Figure S1B, C**). We identified three cell clusters (S11-S13) at the onset of proliferation (**Figure S1A-C**). Most of the upregulated genes in these clusters (35 out of 39) overlapped with those elevated in clusters 4, 5, and 6 from the Seurat analysis (**Figure S1D** and **Table S5**). IPA identified pathways linked to ECM (S11) and cell junction organization (S12 and S13) as active during this phase (**Table S6**).

Both analyses revealed cell groups corresponding to a pre-proliferative state, active proliferation, and a possible shift toward differentiation. The temporal order of these cell groups aligned with a transition from a pre-proliferative state to active proliferation, as indicated by *Mki67*-positive cells at days three and seven. The prominence of cell-cell and cell-ECM interactions, as identified by IPA, was consistent with our prior study, which showed that GER-organoid growth depends on GER cell seeding density at the start of the experiment ^9^. These results defined the transcriptomic profile of GER cells as they initiate proliferation to form growing organoids, which eventually lead to the generation of nascent hair cells and supporting cells. Based on these insights, we hypothesized that a detailed analysis of upregulated genes potentially involved in ECM-modulated growth promotion could provide further mechanistic insight.

### Pharmacological inhibition of galectins and Myc suppresses otic organoid formation and growth

To functionally evaluate candidate pathways associated with the onset of organoid growth, we prioritized genes and pathways based on several criteria, including biological relevance to growth regulation and experimental tractability, rather than differential-expression rank alone. Galectins 1 and 3 were selected because they were upregulated in the onset-associated clusters and represented the prominent cell-cell/ECM signaling signature identified by pathway analysis. Myc was selected as an onset-associated transcriptional regulator with an established role in cell-cycle entry. We additionally tested integrin and enkephalin signaling (**Figure 2D**, **Table S1**, and **Table S3**). Galectins, encoded by *Lgals* genes, are soluble proteins that bind to β-galactoside sugars on glycosylated proteins. They are present in the extracellular matrix but also serve intracellular functions ^19^. Specifically, mRNAs encoding galectin-1 (*Lgals1*) and galectin-3 (*Lgals3*) were differentially upregulated in clusters 4, 5, and 6 compared to all other clusters (**Figure 2D**, **Table S1**, and **Table S3**). Immunofluorescence staining validated galectin-1 expression along the outer surface cell layers of day three organoids, whereas galectin-3 expression was more dispersed (**Figure 3A and B**). Galectin-1 and galectin-3 immunoreactivity was not restricted to Ki67-positive cells, indicating that galectin expression itself does not simply mark the actively cycling population.

**Figure 3.**
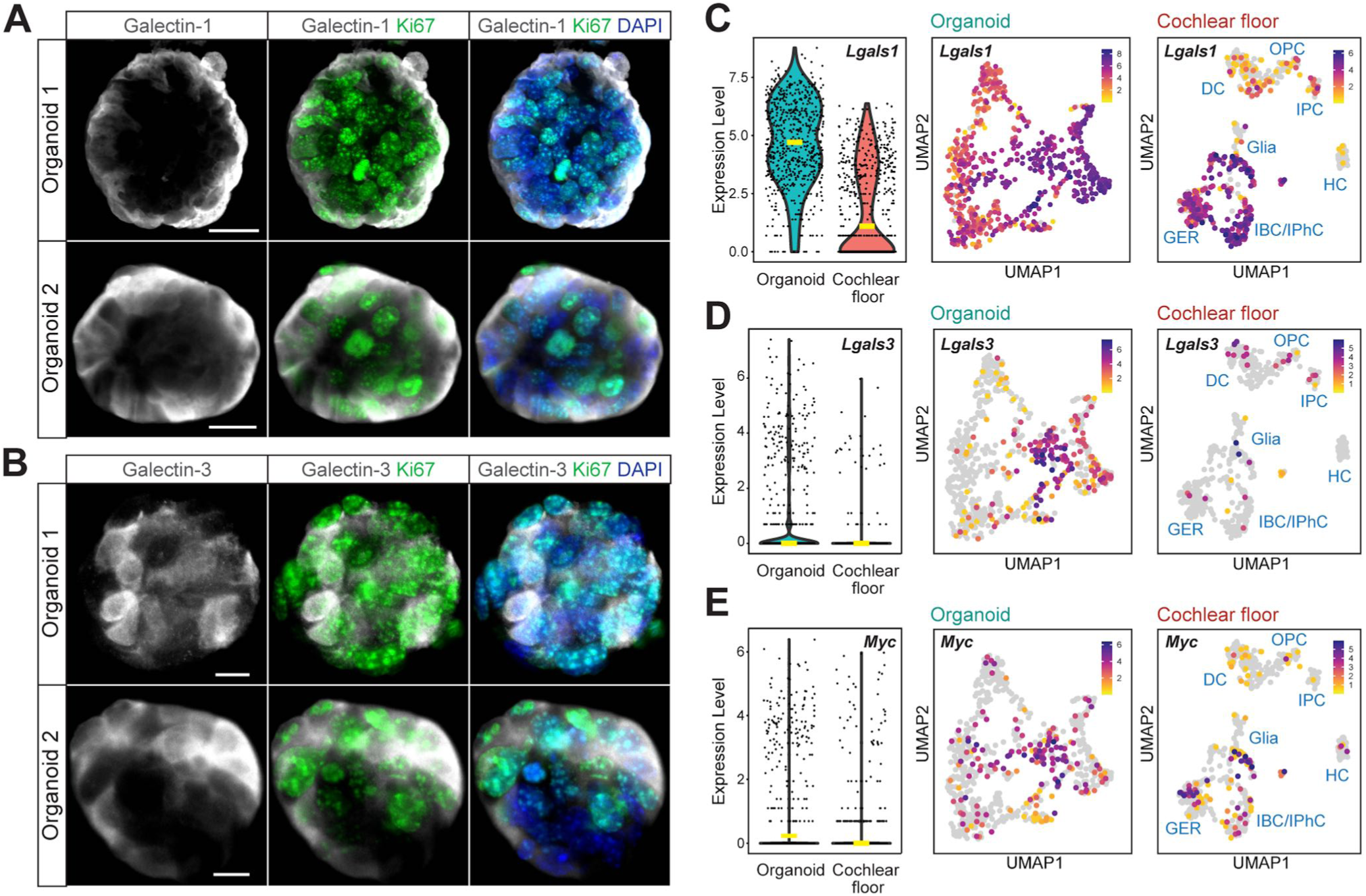
Expression of galectin-1, galectin-3, and Myc in organoids. **(A, B)** Day 3 organoids generated from whole cochlear duct cells. Two representative organoids are shown for each labeling. (A) Immunofluorescence labeling for galectin-1 (grey), Ki67 (green), and nuclei (DAPI, blue). Scale bars: 20 µm. (B) Immunofluorescence for galectin-3 (grey), Ki67 (green), and nuclei (DAPI, blue). Scale bars: 10 µm. (**C-E**) Single-cell RNA-sequencing data analysis comparing expression of galectin-1 (C), galectin-3 (D), and Myc (E) between organoids and the P2 cochlear floor. The organoid UMAPs use the same embedding shown in Figure 2; cluster and D1/D3/D7 timepoint identities are shown in Figure 2A,B. (Left) Violin plots. Yellow bars: median lines. (Center) mRNA expression levels projected on the UMAP plots for GER-organoid cells and (right) for cochlear floor cells. DC: Deiters’ cells; GER: greater epithelial ridge; IBC: inner border cells; IPhC: inner phalangeal cells; OPC: outer pillar cells; IPC: inner pillar cells; HC: hair cells.

We were curious whether galectins were also expressed in P2 cochlear floor cells and analyzed the corresponding single-cell RNA sequencing data ^9^ (**Figure S2** and **Table S7**). In the native P2 cochlear floor dataset, *Lgals1* transcripts were detected prominently in GER and adjacent inner-border/inner-phalangeal populations, whereas *Lgals3* expression was low (**Figure 3C and D**). *Myc* mRNA was detectable in some P2 GER and cochlear floor cells (**Figure 3E**). In organoids, Myc expression was higher and most noticeable in day three organoids, especially in cluster 4 cells, which also displayed the highest levels of *Lgals1* and *Lgals3* mRNA (**Figure 2A and C** and **Figure 3C-E**).

To functionally characterize growth-promoting signals in GER-organoids at the onset of proliferation, we used inhibitors and activators targeting proteins encoded by the genes identified in Seurat clusters 4, 5, and 6 (**Figure 2D**). We used dissociated cochlear duct cells rather than FACS-purified GER cells because our previous work showed that within the cochlear duct, GER cells are the principal cell population capable of organoid formation in media supplemented with growth factors (epidermal growth factor (EGF), fibroblast growth factor 2 (FGF2), insulin growth factor 1 (IGF-1)), and the glycogen synthase kinase 3 (GSK-3) inhibitor CHIR99021 ^9^. Moreover, FACS purification of GER cells from *Sox2^GFP/+^; Fgfr3^CreERT^*^2^*^/+^; Ai14^tdTomato/+^* transgenic mice yields only enough cells to fill a few wells of a 96-well plate, which is insufficient for multi-condition pharmacological screening in organoids.

We selected the IGF-1 receptor inhibitor picropodophyllin ^20^ as a positive control to block IGF-1 and IGF-2 signaling. *Igf2* was differentially expressed at the onset of proliferation (**Figure 2D, Table S1**), and IGF-1 is known to play a critical role in inner ear development and maintenance ^21–24^. Additionally, IGF-1 has been consistently used in otic organoid culture systems ^5,25,26^. We hypothesized that the picropodophyllin inhibition would block IGF-1 supplemented with the culture media, as well as IGF-2 produced by organoid cells at the onset of proliferation. As predicted, picropodophyllin substantially reduced organoid formation (**Figure S3A**).

Next, we targeted galectin-1 and galectin-3 with the selective inhibitors OTX008 and GB1107. Organoid numbers were significantly decreased at 40 µM and 5 µM with OTX008 and GB1107, respectively, while the organoid sizes decreased at 10 µM and 20 µM with the respective inhibitors, as quantified after 7 culture days (**Figure 4A, B, D, and E**). This indicates that although organoid formation is initiated at lower concentrations of OTX008 and GB1107, organoid size and viable cell mass were reduced, indicating impaired organoid growth. These endpoint measurements do not distinguish reduced cell proliferation from impaired cell survival.

**Figure 4.**
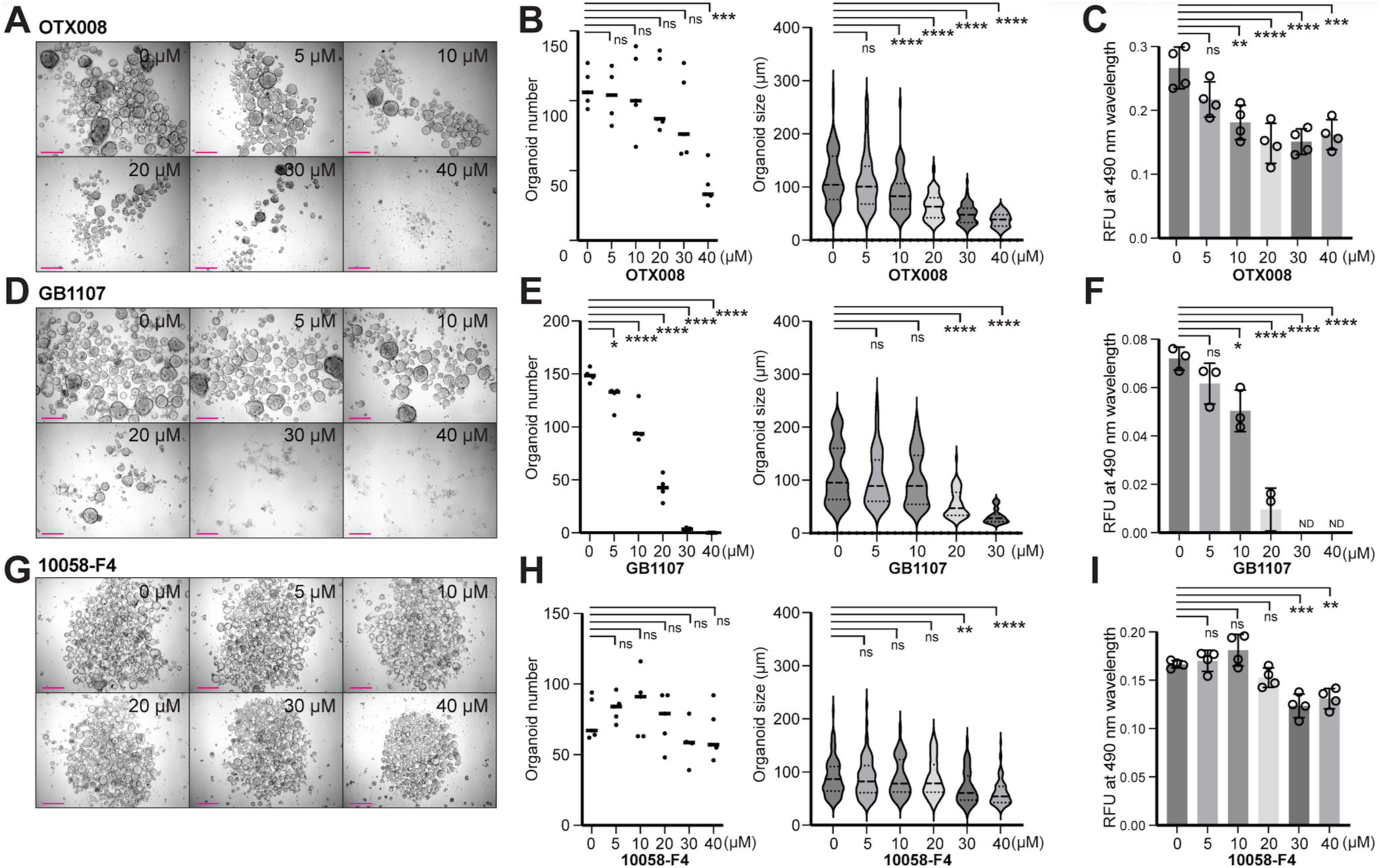
Inhibition of galectin-1, galectin-3, and Myc reduces organoid growth. **(A, D, G)** Dose-dependent inhibition of organoid formation by OTX008 (galectin-1 inhibitor) (A), GB1107 (galectin-3 inhibitor) (D), and 10058-F4 (Myc inhibitor) (G). Organoids generated from whole cochlear duct cells of P2 FVB/NJ mice after 7 days in culture in the presence of increasing inhibitor concentrations. Scale bar: 200 µm. **(B, E, H)** Organoid number (left) and size (right) quantification after 7 days in culture at serial concentrations of inhibitors (OTX008 (B), GB1107 (E), and 10058-F4 (H)). Organoids were centered in the field of view of an inverted microscope in 48-well plates, and quantification was performed using acquired images. For organoid number quantification, medians are shown as solid lines. For organoid size, medians and quartiles are shown as dotted lines. Statistical analysis was performed using Dunnett’s multiple comparisons test in one-way ANOVA. \**p* <0.05, \*\**p* <0.01, \*\*\**p* <0.001, **** *p* <0.0001, ns: not significant. **(C, F, I)** Live cell quantification of organoids using a colorimetric viable cell mass assay after 7 days in culture. Shown are the means ± SD of N = 3-4 independent experiments. Statistical analysis was performed using Dunnett’s multiple comparisons test in one-way ANOVA. \**p* <0.05, \*\**p* <0.01, \*\*\**p* <0.001, **** *p* <0.0001, ns: not significant.

Because organoids vary in morphology from densely cellular to those with an internal acellular cavity, and they are not always spherically shaped, the organoid numbers and sizes do not necessarily reflect the resulting number of cells after proliferation. We therefore used a colorimetric assay as an orthogonal measure of metabolically active (viable) cell mass. The viable cell mass assay showed a dose-dependent decrease with both OTX008 and GB1107, with a significant effect observed at 10 µM for each (**Figure 4C and F**). Inhibition of Myc with 10058-F4 was modest yet clearly detectable in organoid size at 30 µM, while organoid number stayed consistent regardless of the drug concentration (**Figure 4G and H**). The viable cell mass assay showed a significant decrease at 30 µM (**Figure 4I**).

To confirm that these selective inhibitors affect GER cell-derived organoids, we also tested them with FACS-enriched GER cells. Organoid number was zero at 20 µM, 10 µM, and 40 µM with OTX008, GB1107, and 10058-F4, respectively (**Figure 5A**). Because these assays were initiated with only 2,000 FACS-enriched GER cells per well and absolute organoid numbers and sizes varied between independent sorting experiments, organoid number was used qualitatively in this experiment. Quantitative dose-response effects were assessed using the viable cell mass assay, which measures the entire well. The viable cell mass assay showed a dose-dependent reduction in organoid growth, with significant inhibition at 5 µM for both OTX008 and GB1107, and at 40 µM with 10058-F4 (**Figure 5B**). Our results show that pharmacological inhibition of galectin-1, galectin-3, or Myc suppresses GER-derived organoid formation and reduces viable cell mass. Because these endpoint assays cannot distinguish impaired cell-cycle progression from reduced cell survival, we interpret them as evidence that these pathways contribute to organoid formation and growth.

**Figure 5.**
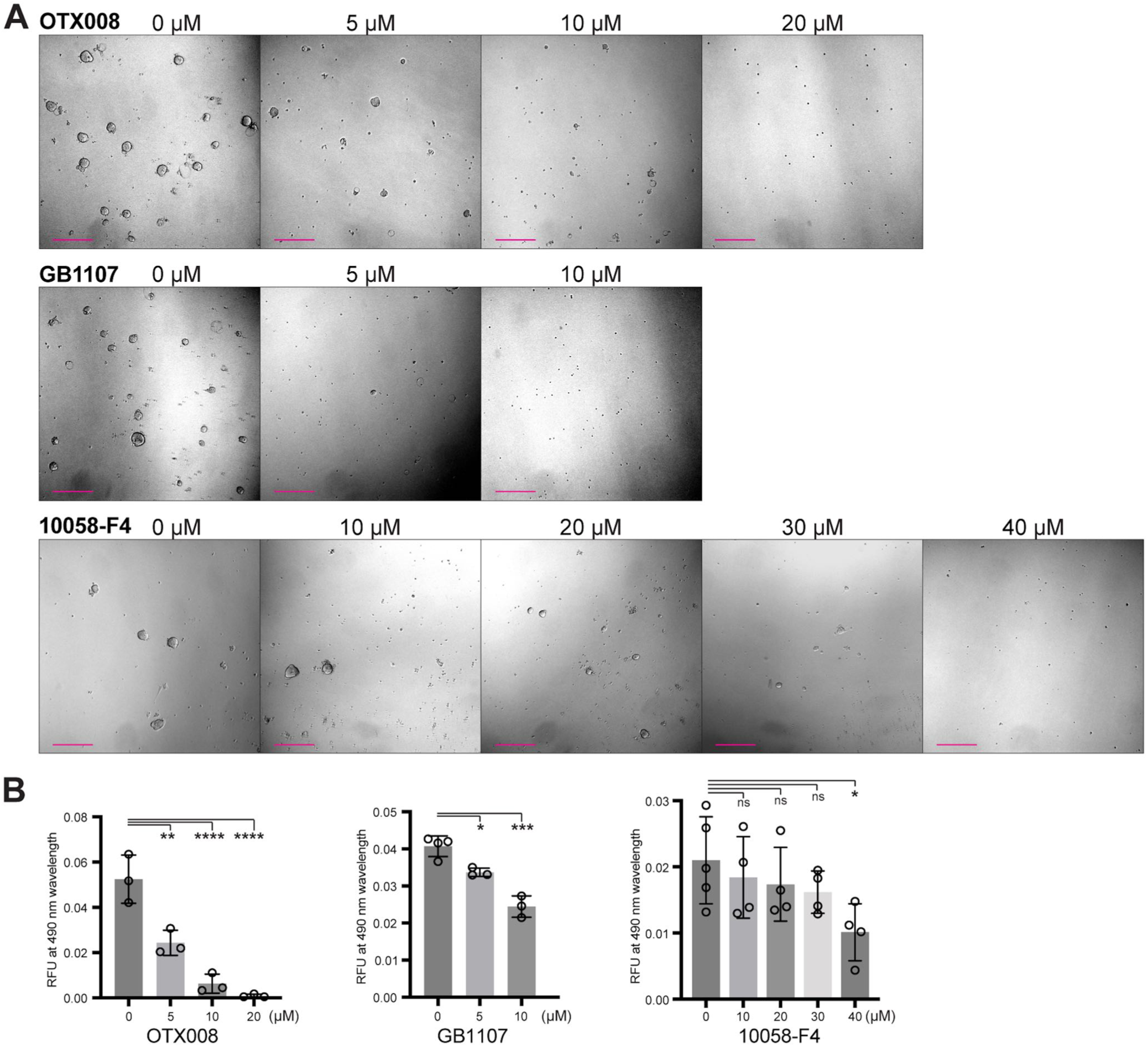
GER cell-derived organoid growth is strongly attenuated with inhibitors targeting galectin-1, galectin-3, and Myc. **(A)** Representative images of organoids generated from FACS-enriched GER cells after seven days in culture, illustrating the dose-dependent reduction in detectable organoid formation following treatment with OTX008 (galectin-1 inhibitor), GB1107 (galectin-3 inhibitor), and 10058-F4 (Myc inhibitor). Scale bars: 200 µm. **(B)** Quantification of viable cell mass using a colorimetric live cell assay after seven days in culture in increasing concentrations of OTX008, GB1107, and 10058-F4. Means ± SDs of N = 3-4 independent experiments are shown. Statistical analysis was performed using Dunnett’s multiple comparisons test in one-way ANOVA. \**p* <0.1, \*\**p* <0.01, \*\*\**p* <0.001, **** *p* <0.0001, ns: not significant.

A limitation of our pharmacological screening strategy is that not all signaling proteins identified by differential gene expression analysis have specific chemical compound modulators. We partially addressed this constraint with compounds that broadly inhibit signaling mediated through integrins. Our analysis identified a substantial number of genes encoding ECM components that directly interact with integrins, such as collagens (*Col18a*, *Col1a*, *Col2a*, *Col5a*, *Col6a*, *Col9a*), tenascin C (*Tnc*), laminin (*Lamc2*), urokinase receptor (*Plaur*), along with proteins that indirectly form complexes with integrins (**Figure 2D**, **Table S2**, and **Table S3**) ^27–29^. Integrins serve as linkers between the ECM and the cytoskeleton, interact with a wide range of proteins, and transduce signals into cells ^28^. We utilized three potent inhibitors (RGDS peptide, echistatin, and ATN-161) that target different sets of integrin heterodimer subunits; however, none of them significantly inhibited organoid growth (**Figure S3B**).

Based on the observed upregulation of *penk*, which encodes the preprotein for the endogenous opioid enkephalins (**Figure 2D** and **Table S3**), we tested enkephalins as potential activators. In addition, we applied Naltrexone as an opioid receptor antagonist ^30^. These modulators did not induce significant changes in organoid formation (**Figure S3C**).

Together, these pharmacological experiments identify galectin and Myc signaling as candidate regulators of organoid formation and growth, leaving the functional relationship between these pathways to be determined.

### Overexpression of galectin-1 and Myc increases otic organoid growth

Next, we asked whether overexpression of galectin-1, galectin-3, or Myc could enhance organoid formation and growth. Adeno-associated virus (AAV)-mediated overexpression of genes was conducted in cochlear duct cells isolated from P2 mice. We used dissociated cochlear duct cells for the assay rather than enriched GER cells because AAV infection prevented organoid formation from FACS-enriched GER cells, likely due to limited starting cell numbers, regardless of the encoded genes. GER cells are the principal organoid-forming cell group within dissociated cochlear duct at P2^9^, although we cannot completely exclude contribution from non-GER cochlear epithelial cell populations to the resulting organoids. We first assessed the ability of AAVs to transduce organoid cells using a control AAV serotype DJ encoding the EGFP reporter gene driven by the CAG promoter (AAV-EGFP, **Figure S4A**). At 3 days post-infection (3 dpi), organoids transduced with AAV-EGFP exhibited robust EGFP expression, while no EGFP expression was observed in uninfected negative controls (**Figure S4B**). We quantified the number of AAV-EGFP-transduced cells with detectable fluorescence through FACS analysis and determined that, on average, 84% of organoid cells expressed EGFP (**Figure S4C**).

For functional studies, we used AAVs encoding *Lgals1* (AAV-lgals1), *Lgals3* (AAV-lgals3), and *Myc* (AAV-myc) (**Figure S4A**). We chose a 9 dpi endpoint because AAV-mediated gene expression takes up to 48 hours, and seven additional days of organoid growth would provide sufficient time to compare growth differences (**Figure 6A**). To evaluate cell proliferation, we conducted quantitative PCR (qPCR) for *Mki67* mRNA at 3, 5, 7, and 9 dpi. Both AAV-lgals1 and AAV-myc transduction resulted in increased mRNA levels of the proliferation marker *Mki67*, compared to AAV-EGFP control-transduced organoids. AAV-lgals1 induced a gradual increase in *Mki67* over time, while AAV-myc transduction triggered a rapid and more pronounced rise in *Mki67* mRNA levels at 3 dpi (**Figure 6B**). Consistent with increased *Mki67*, *Lgals1* mRNA gradually increased in AAV-lgals1-transduced organoids and reached approximately 8-fold higher levels at 9 dpi, compared to AAV-EGFP-transduced organoids. *Myc* mRNA overexpression was effective at 3 dpi, reaching levels 15-fold higher, subsequently decreasing to approximately 10-fold at 5-9 dpi (**Figure 6C**). Although both vectors were driven by the CAG promoter, their expression cassettes differed, and qPCR measured total steady-state target mRNA rather than promoter activity. We therefore interpret these measurements as demonstrating effective but distinct temporal profiles of transgene expression rather than directly comparing the relative potency of *Lgals1* and *Myc* in promoting organoid growth. Overexpression of *Lgals1* significantly increased organoid growth compared to the control, as quantified by the number of organoids and the viable cell mass assay at 9 dpi. Also AAV-Myc significantly increased organoid number; however, viable cell mass at 9 dpi was not significantly different from the AAV-EGFP control (**Figure 6D-F**). To determine whether differences in viable cell mass were associated with cell death, we dissociated organoids at 9 dpi, labeled dead cells with Far Red Dead Cell Stain, and quantified the proportion of viable cells by flow cytometry (**Figure 6G**). AAV-Myc-treated cultures contained a significantly lower proportion of viable cells than control or AAV-Lgals1-treated cultures. At 9 dpi, approximately 77% of AAV-Myc-treated cells remained viable, compared with >90% in the control and AAV-Lgals1 cultures. These results indicated that Myc induced a strong early proliferative response, while prolonged overexpression was accompanied by increased cell death. Because there was no significant change in the live cell proportion at 9 dpi in galectin-1-overexpressing organoids compared to the control, we conclude that the increase in organoid growth results from enhanced cell proliferation (**Figure 6G**). Unlike galectin-1 and Myc, galectin-3 overexpression did not significantly affect organoid growth compared to the control (**Figure 6H-J**). Together, these experiments show that AAV-mediated overexpression of galectin-1 or Myc enhances organoid formation and/or growth in neonatal cochlear duct cultures. Myc additionally induced an early increase in Mki67 expression, although prolonged Myc overexpression was accompanied by reduced cell survival.

**Figure 6.**
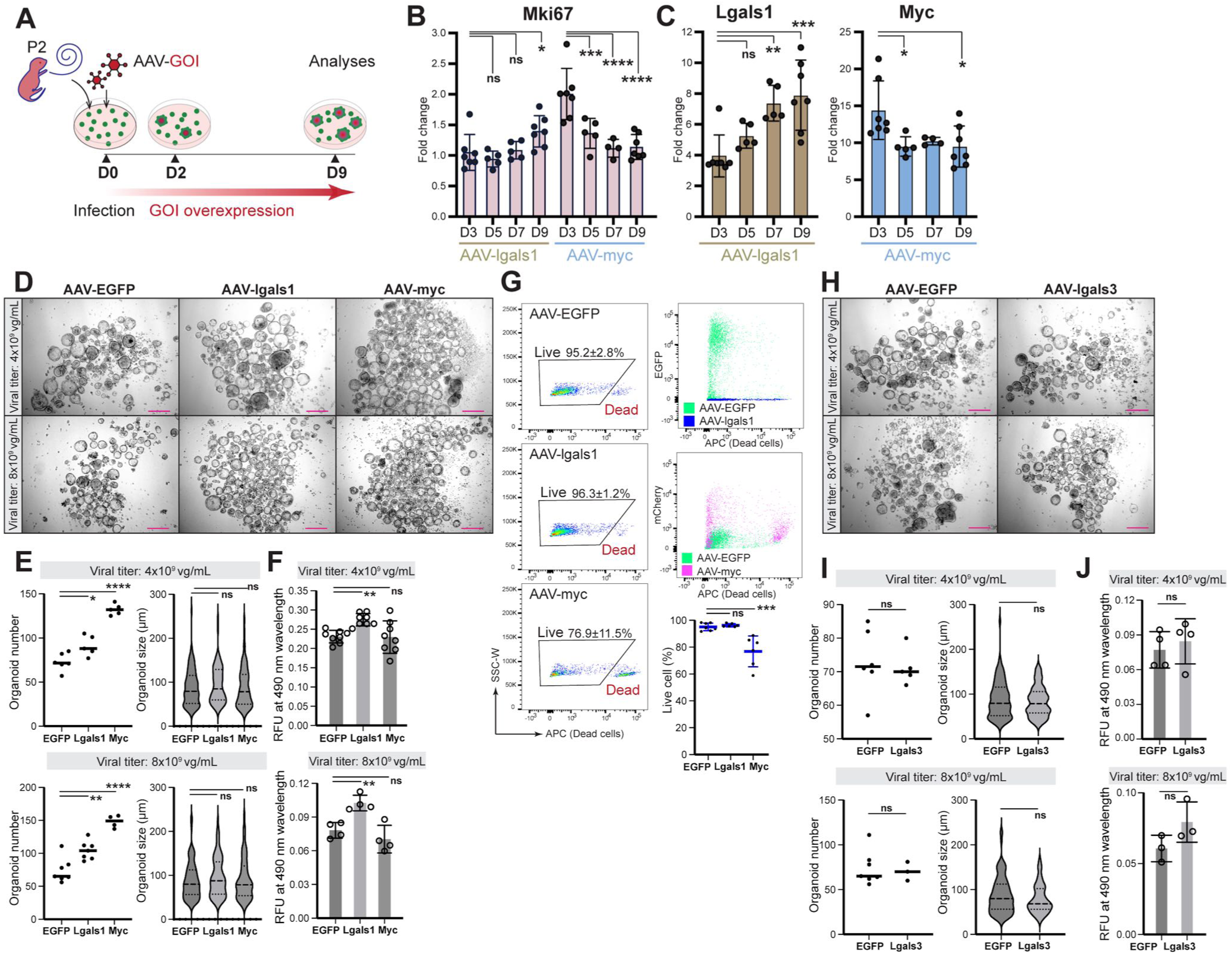
AAV-mediated overexpression of galectin-1, galectin-3, and Myc in organoids generated from neonatal cochlear duct cells. **(A)** Schematic illustration of the organoid formation assay using dissociated P2 FVB/NJ mouse cochlear duct cells infected with AAVs for each gene of interest (AAV-GOI). Effective gene overexpression was presumed to start within two days post-infection (dpi), and cell proliferation was assessed at 9 dpi. **(B)** qPCR for *Mki67* mRNA in organoids infected with AAV-lgals1 or AAV-myc, compared to organoids infected with AAV-EGFP. Statistical analysis was performed using Dunnett’s multiple comparisons test in one-way ANOVA. \**p* <0.05, \*\*\**p* <0.001, \*\*\*\**p* <0.0001, ns: not significant. **(C)** mRNA expression changes for *Lgals1* and *Myc* were quantified using qPCR in organoids infected with AAV-lgals1 (left) and AAV-myc (right) at 3, 5, 7, and 9 dpi, compared to organoids infected with AAV-EGFP for each respective time point. Statistical analysis was performed using Dunnett’s multiple comparisons test in one-way ANOVA. \**p* <0.05, \*\**p* <0.01, \*\*\**p* <0.001, ns: not significant. **(D)** Cochlear duct cells were infected with AAV-EGFP, AAV-lgals1, or AAV-myc at titers of 4x10^9^ or 8x10^9^ viral genomes per milliliter (vg/mL). AAV-lgals1 and AAV-myc visibly enhanced organoid formation compared to the control (AAV-EGFP) at both viral titers at 9 dpi. Scale bar: 200 µm. **(E)** Quantification of organoid number (left) and size (right) at 9 dpi following infection with AAV-EGFP, AAV-lgals1, or AAV-myc at 4x10^9^ or 8x10^9^ vg/mL. Organoids were centered in the field of view of an inverted microscope in 48-well plates, and quantification was performed using acquired images. For organoid number quantification, medians are shown as solid lines. For organoid size, medians and quartiles are shown as dotted lines. Statistical analysis was performed using Dunnett’s multiple comparisons test in one-way ANOVA. \**p* <0.05, \*\**p* <0.01, **** *p* <0.0001, ns: not significant. **(F)** Live cell quantification of the organoids using a colorimetric viable cell mass assay. Means ± SDs are shown. N = 8 and N = 4 for the experiments with 4x10^9^ and 8x10^9^ vg/mL viral titers. Statistical analysis was performed using Dunnett’s multiple comparisons test in one-way ANOVA. \**p* <0.05, \*\**p* <0.01, ns: not significant. **(G)** FACS quantification of live cells in organoids at 9 dpi following infection with AAV-EGFP, AAV-lgals1, or AAV-myc at 4x10^9^ vg/mL. (Left three panels) Individual live cell populations lacking dead cell stain cells (detected in the allophycocyanin (APC) channel), are gated in dot plots. (Right top two panels) Cells from AAV-EGFP- and AAV-lgals1-infected organoids, as well as cells from AAV-EGFP- and AAV-myc-infected organoids, are overlaid to show the expression of EGFP and mCherry encoded in AAV-EGFP and AAV-myc, respectively. (Right bottom panel) Percentage of live cells in organoids at 9 dpi following infection with AAV-EGFP, AAV-lgals1, or AAV-myc at 4x10^9^ vg/mL. Statistical analysis was performed using Dunnett’s multiple comparisons test in one-way ANOVA. \*\*\**p* <0.001, ns: not significant. **(H)** Cochlear duct cells were infected with AAV-EGFP or AAV-lgals3 at titers of 4x10^9^ or 8x10^9^ vg/mL. AAV-lgals3-infected organoids exhibited no significant difference in growth compared to the control (AAV-EGFP) at both viral titers at 9 dpi. Scale bar: 200 µm. **(I)** Quantification of organoid number (left) and size (right) at 9 dpi following infection with AAVEGFP, or AAV-lgals3 at 4x10^9^ or 8x10^9^ vg/mL. Organoids were centered in the field of view of an inverted microscope in 48-well plates, and quantification was performed using acquired images. For organoid number quantification, medians are shown as solid lines. For organoid size, medians and quartiles are shown as dotted lines. Statistical analysis was performed using the two-tailed Mann-Whitney U test. ns: not significant. **(J)** Live cell quantification of organoids using a colorimetric viable cell mass assay. Means ± SDs of N = 3-4 experiments are shown. Statistical analysis was performed using a two-tailed, unpaired t-test. ns: not significant.

### Myc, but not galectin-1, promotes organoid growth from third-week cochlear cells

Otic organoid formation is a characteristic of the neonatal cochlea, particularly of GER cells, and is observed only before the onset of hearing in mice, which occurs around P12 ^31^. Since AAV-mediated overexpression of *Lgals1* and *Myc* mRNA enhanced organoid growth at P2, we next investigated whether overexpression of galectin-1 and Myc would stimulate organoid growth from post-neonatal (P14-P17) cochlear cells. To purify cochlear epithelial cells, we utilized the epithelial cellular adhesion molecule (EpCAM, CD326), which is expressed in the cochlear floor and adjacent epithelia (**Figure 7A**). At this developmental stage, the GER has already regressed (**Figure 7A**), and active cell proliferation is absent, resulting in markedly reduced capacity for organoid formation ^32^. As antibodies to EpCAM have previously been used to enrich cochlear epithelial cells ^33^, we applied magnetic cell separation (MACS) with an anti-CD326 (EpCAM) antibody to isolate cells from P14-P17 mouse cochleae (**Figure 7B**).

**Figure 7.**
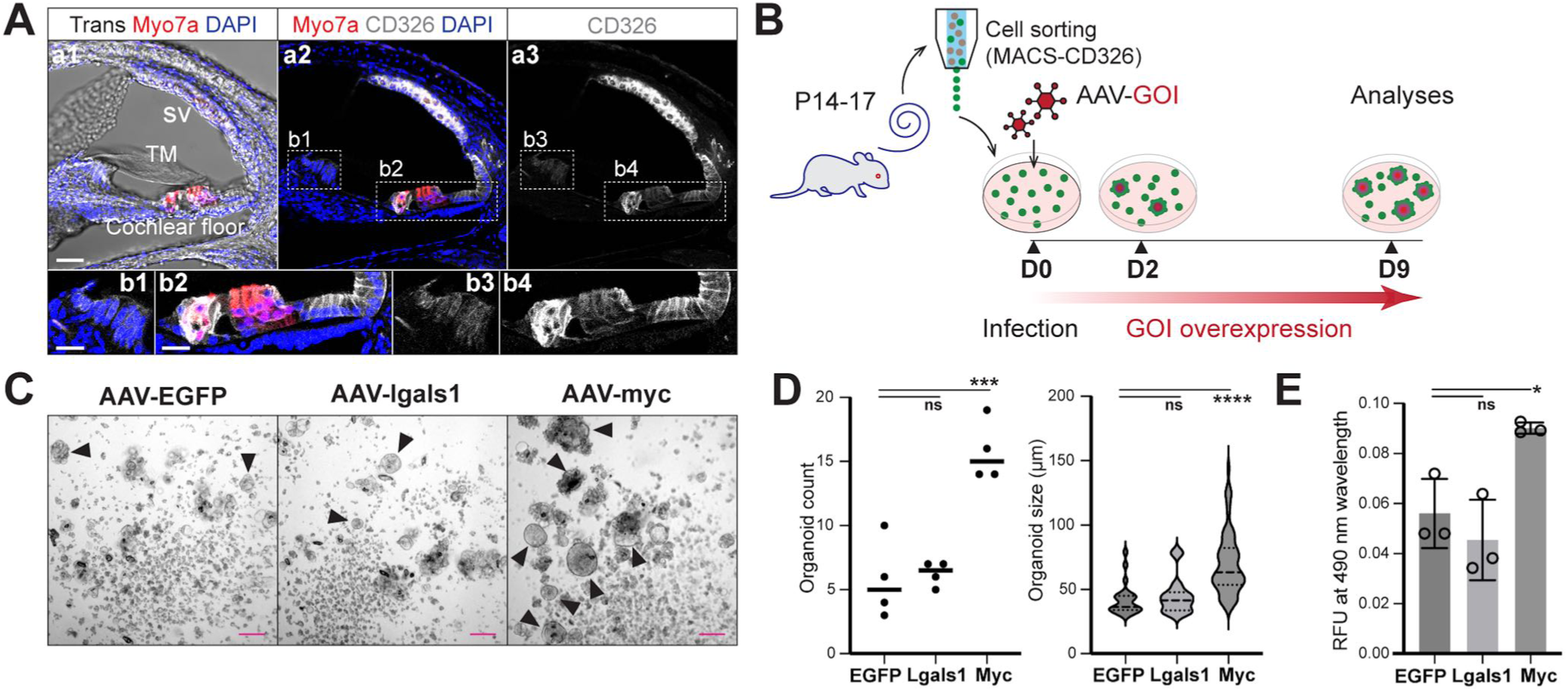
Overexpression of galectin-1 and Myc in organoids generated from postnatal day 14-17 cochlear epithelial cells. **(A)** (a1-3) Immunofluorescence detection of Myosin 7a (Myo7a) and CD326/EpCAM in a P14 FVB/NJ mouse cochlea vibratome section. Hair cells are labeled for Myosin 7a. CD326/EpCAM was detected in cochlear floor epithelial cells and the cells of the stria vascularis. The GER was absent at this more mature age. SV: stria vascularis; TM: tectorial membrane. Scale bar: 50 µm. (b1-4) Magnified images of the areas surrounded by the dotted boxes in a2 (b1,2) and a3 (b3,4). Scale bar: 25 µm. **(B)** Schematic illustration of the organoid formation assay timeline using dissociated P14-17 FVB/NJ mouse cochlear duct cells infected with AAVs for each gene of interest (AAV-GOI). CD326-positive epithelial cells were enriched by Magnetic Activated Cell Sorting (MACS). Organoids were infected with AAV-GOIs. Gene overexpression was presumed to start within two days post-infection, and cell proliferation was assessed at 9 dpi. **(C)** MACS-enriched CD326-positive cochlear cells were infected with AAV-EGFP, AAV-lgals1, or AAV-myc at a titer of 4x10^9^ vg/mL. AAV-myc infection visibly enhanced organoid formation, while AAV-lgals1-infected organoids exhibited no significant difference in growth compared to the control (AAV-EGFP). Arrowheads: organoids. The cellular debris likely represents the remnants of cochlear cells that did not grow into organoids at 9 dpi. Scale bar: 100 µm. **(D)** Quantification of organoid number (left) and size (right) at 9 dpi following infection with AAV-EGFP, AAV-lgals1 or AAV-myc at 4x10^9^ vg/mL. Organoids were centered in the field of view of an inverted microscope in 48-well plates, and quantification was performed using acquired images. For organoid number quantification, medians are shown as solid lines. For organoid size, medians and quartiles are shown as dotted lines. Statistical analysis was performed using Dunnett’s multiple comparisons test in one-way ANOVA. *** *p* <0.001, **** *p* <0.0001, ns: not significant. **(E)** Live cell quantification of the organoids using a colorimetric viable cell mass assay. Means ± SDs of three experiments are shown. Statistical analysis was performed using Dunnett’s multiple comparisons test in one-way ANOVA. \**p* <0.05, ns: not significant.

The purified cells were then transduced with AAVs for *Lgals1* and *Myc* mRNA overexpression, as well as with a control virus (AAV-EGFP). Cells infected with AAV-lgals1 exhibited low levels of organoid formation that did not differ from the control. In contrast, cells infected with AAV-myc displayed significantly increased organoid formation, as reflected in the number, size, and live-cell mass (**Figure 7C-E**). These results suggest that galectin-1 alone primarily enhances cell proliferation of neonatal GER cells, whereas Myc can promote proliferation akin to P2 GER cells, resulting in increased organoid formation from the EpCAM-enriched P14-P17 cochlear cell population.

### Galectin-1 is upregulated in the GER after cochlear supporting cell ablation

To explore the *in vivo* relevance of our findings, we employed an Lgr5-DTR mouse model in which the diphtheria toxin (DT) receptor is driven by endogenous Leucine-rich repeat containing G protein-coupled receptor 5 (Lgr5) expression ^34^. Previous studies have demonstrated that intraperitoneal (IP) injection of DT in P1 mice induces Lgr5-driven cell ablation, resulting in the loss of inner phalangeal cells (IPhCs), inner border cells (IBCs), the third row of Deiters’ cells, and inner pillar cells at P3-P4. Following DT-mediated Lgr5+ cell ablation, damage-induced proliferation is robust at P3-P4, particularly in the lateral GER of the apical and middle cochlear turns, followed by the emergence of new IPhCs at P7 ^8,12^.

We investigated gene expression changes in GER cells after DT-mediated organ of Corti cell ablation at P1 by analyzing single-cell RNA sequencing data obtained from P4 Lgr5-DTR mice (damage model: *Ki67^CreERT^*^2^*^/+^; Lgr5^DTR/+^; Rosa26R^tdTomato/+^*) and control mice (control: *Ki67^CreERT^*^2^*^/+^; Rosa26R^tdTomato/+^*) (**Figure 8A**) ^8,12,34^. Distinct cell clusters, including medial GER, lateral GER, IPhCs/IBCs, and proliferating cells, were identified based on the expression of specific known marker genes (**Figure 8B, C, E**, and **Table S8**). The cells expressing proliferation markers Mki67 or Top2a were almost exclusively detected in the damage model in cluster 8 (**Figure 8B, C, E, F**, arrows in **E**). This cluster was inferred to consist of lateral GER cells that entered the cell cycle in response to damage. As expected, we noted lower levels of *Lgr5* mRNA in the damage model compared to the control (**Figure 8D**). Moreover, supporting cell markers that are also expressed in the lateral GER, such as *Sox2*, *Jag1*, and *Fabp7* ^9,35^, were downregulated in the damage model, while proliferation markers, including *Cdk4* and *Ccnd1* ^36^, were upregulated (**Figure 8H** and **Table S8**) compared to the control. These observations confirmed that Lgr5-driven cell loss, along with active cell proliferation, occurs at P4 in the damaged cochleae. In the control mice, *Lgals1* was highly expressed in the medial GER, and its expression was further increased in the damage model (**Figure 8G, H**, and **Table S8**). *Myc* expression was also upregulated in the medial GER in the damage model (**Figure 8F, G**, and **Table S8**). Upregulation of Galectin-1 protein was validated in Lgr5-DTR (*Lgr5^DTR/+^*) mice at P4 in the GER following DT treatment at P1, compared with controls (**Figure 8I**). Notably, there was prominent expansion of the galectin-1 expression domain from the medial to lateral GER in the damaged cochleae, accompanied by the loss of fatty acid binding protein 7 (Fabp7)-expressing cells, including IPhCs and IBCs, and the emergence of proliferating cells in lateral GER, marked by Ki67 (**Figure 8I**).

**Figure 8.**
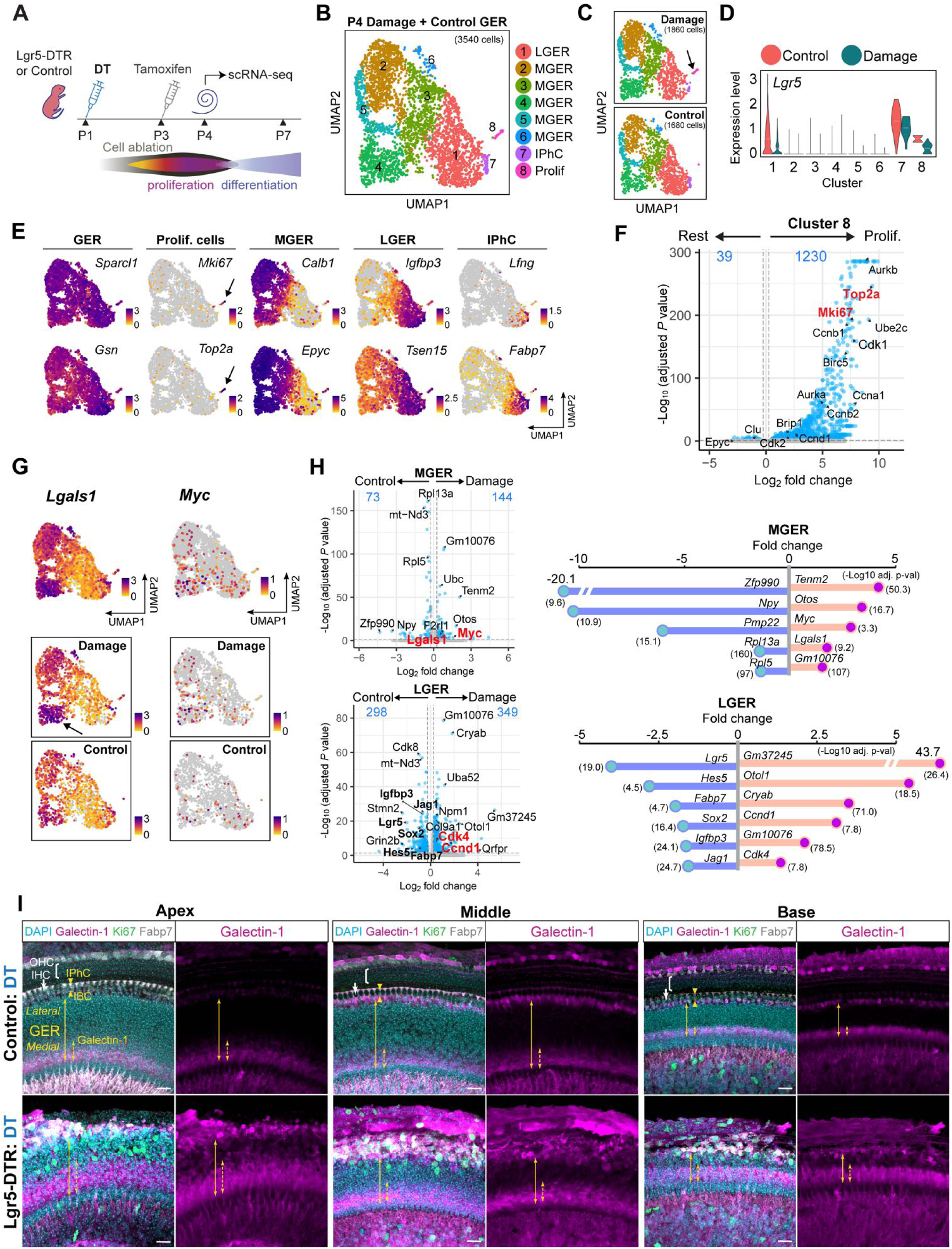
Galectin-1 upregulation in the GER in the Lgr5-DTR mouse damage model. **(A)** Schematic illustration of the Lgr5-DTR mouse damage model. Intraperitoneal injection of diphtheria toxin (DT) in P1 mice induces Lgr5-driven cell ablation. Cell loss and concurrent cell proliferation in the lateral GER occur by P4. New IPhCs emerge by P7. **(B)** Seurat data analysis with 3,773 cells obtained from P4 Lgr5-DTR damage model mice (*Ki67^CreERT2/+^; Lgr5^DTR/+^; Rosa26R^tdTomato/+^*) and control (*Ki67^CreERT2/+^; Rosa26R^tdTomato/+^*). Eight clusters representing medial GER (MGER), lateral GER (LGER), IPhCs (this cluster also includes IBCs), and proliferating cells (Prolif) are visualized in a UMAP projection. **(C)** UMAP visualization of the clusters split by damage and control mouse models. Cluster 8 (arrow) is predominantly composed of cells after damage. **(D)** Violin plot showing the downregulation of Lgr5 expression in the damage model compared to the control. **(E)** Marker-gene expression used for cell-type identification, projected onto the integrated damage + control UMAP: GER (*Sparcl1* and *Gsn*), proliferating cells (*Mki67* and *Top2a*), MGER (*Calb1* and *Epyc*), LGER (*Igfbp3* and *Tsen15*), and IPhC (*Lfng* and *Fabp7*). Condition-specific expression differences are shown in panels G and H. A Log2 expression scale is shown for each UMAP. **(F)** Volcano plot showing differential gene expression in cluster 8 (proliferating cells) compared to the remaining clusters. **(G)** *Lgals1* (left) and *Myc* (right) expression, projected onto the UMAP plot, and visualized individually for the damage model and control cells. The arrow points to cells with high expression of *Lgals1*. **(H)** Differentially expressed genes were analyzed in cells from the damage model compared to the control, individually for MGER (clusters 2, 3, 4, 5, and 6) (upper) and LGER (cluster 1) (lower) clusters. Genes highlighted in red (*Cdk4*, *Ccnd1*, *Myc*, and *Lgals1*) are proliferation-associated genes. The fold-change expression of selected genes is shown with lollipop plots for MGER and LGER clusters. **(I)** Immunohistology of P4 cochleae from control and Lgr5-DTR mice (*Lgr5^DTR/+^*) post-DT treatment revealed upregulation of galectin-1 in the medial and lateral GER. Proliferating cells labeled with Ki67 were observed in the lateral GER, accompanied by a loss of Fabp7-expressing cells (IPhC and IBC) along the apex-to-base gradient. Solid yellow arrows indicate the GER, while dotted yellow arrows mark areas with visible galectin-1 expression. IHC: inner hair cell; OHC: outer hair cell; IPhC: inner phalangeal cell; IBC: inner border cell. Scale bars: 20 µm.

### *In vivo* inhibition of galectin-1 suppresses GER cell proliferation

To functionally investigate the role of galectin-1 *in vivo* during the regenerative proliferation of GER cells, we administered the inhibitor OTX008 (IP, 10 mg/kg) to Lgr5-DTR mice (*Lgr5^DTR/+^*) and controls at P2 and P3, following DT treatment at P1 (**Figure 9A**). Cochleae were harvested at P4, and cell loss was assessed by counting Fabp7-positive IPhCs and IBCs. DT-treated control cochleae exhibited an organized arrangement of Fabp7-positive cells, with no Ki67-labeled cells observed (**Figures 9B**, and **S5A, B**).

**Figure 9.**
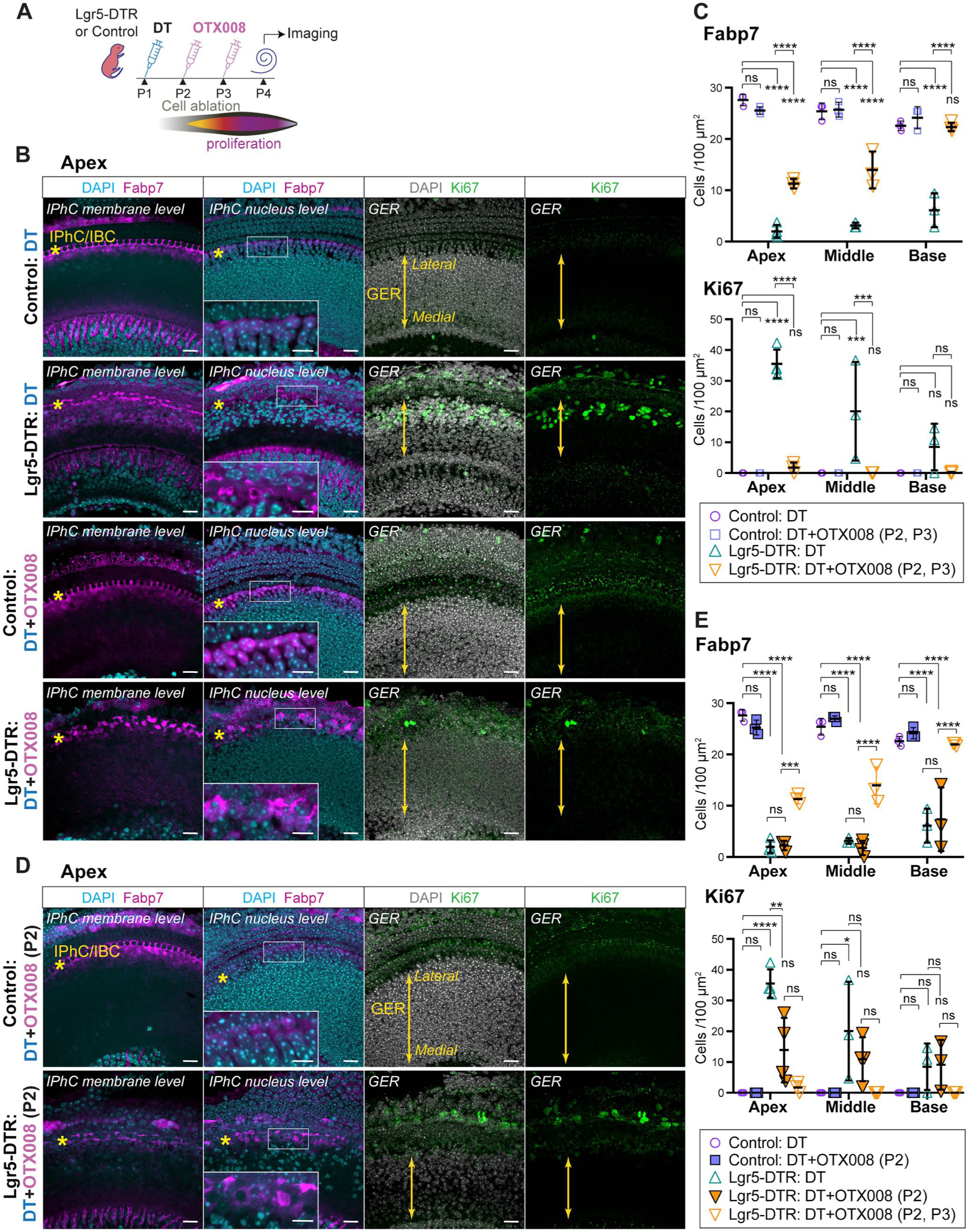
*In vivo* administration of the galectin-1 inhibitor OTX008 prevents GER cell proliferation. **(A)** A schematic illustrating the DT and OTX008 treatment schedule in Lgr5-DTR (*Lgr5^DTR/+^*) and control mice. **(B)** Immunohistology of P4 cochleae obtained from control and Lgr5-DTR mice treated with DT at P1, representing intact and damage controls, respectively, and from control and Lgr5-DTR mice treated with OTX008 (10 mg/kg per injection) at P2 and P3. Stacked images of the apical turns of the cochleae are presented at three levels: IPhC apical membrane, IPhC nuclei, and GER (scale bars: 20 µm). In the IPhC nucleus-level images, magnified views of the IBC/IPhC nuclei are shown for the areas outlined by squares (scale bars: 10 µm). Fabp7-positive debris lacking nuclei, along with pyknotic cells, were observed in Lgr5-DTR cochleae treated with DT (damage control) or DT+OTX008. Asterisks indicate IPhCs and IBCs. Solid yellow arrows indicate the GER. **(C)** Quantification of Fabp7-positive and Ki67-positive cells in the apical, middle, and basal turns of the cochleae, corresponding to Figures 7B and S5A, B. **** *p*<0.0001, *** *p*<0.001, ** *p*<0.01, * *p*<0.05, ns: not significant. **(D)** P4 cochleae obtained from control and Lgr5-DTR mice treated with DT at P1, followed by OTX008 (10 mg/kg per injection) only at P2. Stacked images of the apical turns of the cochleae are presented at three levels, as described in (B). Scale bars: 20 µm and 10 µm (magnified images). Fabp7-positive debris lacking nuclei, along with pyknotic cells, were observed in Lgr5-DTR cochleae treated with DT+OTX008 (P2). Asterisks indicate IPhCs and IBCs. Solid yellow arrows indicate GER. **(E)** Quantification of Fabp7-positive and Ki67-positive cells in the apical, middle, and basal turns of the cochleae, corresponding to Figures 7D and S6A, B. **** *p*<0.0001, *** *p*<0.001, ** *p*<0.01, * *p*<0.05, ns: not significant.

In contrast, in DT-treated Lgr5-DTR cochleae, we observed a decrease of Fabp7-positive cells in the IPhC and IBC region together with a concomitant increase in Ki67-positive cells in the lateral GER along the apex-to-base gradient (**Figures 9B, C,** and **S5A, B**). Neither cell loss nor cell proliferation was noted in DT-treated control mice injected with OTX008, confirming that OTX008 itself does not affect quiescent cochlear cells (**Figures 9B, C,** and **S5A, B**). Lgr5-DTR mice treated with DT and OTX008 exhibited a significant decrease in Fabp7-positive cells in the apical and middle turns of the cochleae compared to controls. The inhibition of galectin-1 signaling resulted in a substantial and significant reduction of Ki67-positive cells, supporting a role for galectin-1 signaling in damage-induced GER cell proliferation (**Figures 9B, C**, and **S5A, B**). The observed effect may reflect either direct inhibition of galectin-1 within the cochlea or an indirect systemic effect of OTX008 administration.

It should be noted that OTX008 administration partially mitigated the Fabp7-positive cell loss compared to DT-treated Lgr5-DTR damage control, with the effect being more pronounced towards the basal turn. The partial preservation of Fabp7-positive cells following OTX008 treatment may reflect context-dependent effects of galectin-1 signaling on proliferation and survival. Because it remained unclear whether the decrease in proliferating cells in the GER resulted from incomplete ablation or was a primary effect of galectin-1 inhibition by OTX008, we further investigated this by administering OTX008 (IP, 10 mg/kg) only at P2 in Lgr5-DTR mice after DT treatment at P1. Although the number of Fabp7-positive cells decreased to an equivalent level as the damage control, a significant reduction in Ki67-positive cells was observed in the GER, showing that reduced damage-induced GER proliferation persisted even when Fabp7-positive cell loss was comparable to the damage control (**Figures 9D, E**, and **S6A, B**). The apex-to-base gradient of damage-induced GER proliferation parallels the developmental gradient of cochlear maturation. Thus, the stronger proliferative response observed in the apical and middle GER may reflect persistence of a less mature, more plastic cellular state.

These findings demonstrate that galectin-1 plays a role in damage-induced cell proliferation in the GER, contributing to the replenishment of lost cells. Importantly, they also suggest that the cell proliferation mechanism in GER-derived organoids parallels the *in vivo* mechanism of GER proliferation.

## Discussion

Certain tissues retain relatively high proliferative and regenerative capacity in neonatal stages in mammals, whereas this capacity is limited or absent in adults. These tissues include the heart, Achilles tendon, brain, spinal cord, and the annulus fibrosus of the intervertebral disc, where extracellular matrix and cytoskeletal remodeling provide a microenvironment that supports regeneration ^37,38^. Cochlear epithelium is one such tissue: the GER (Kölliker’s organ), located adjacent to the modiolar edge of the organ of Corti is replaced by inner sulcus cells during the second and third weeks of postnatal cochlear maturation.

Several lines of evidence suggest that GER cells possess otic progenitor characteristics. First, ectopic expression of Atoh1 in neonatal cochlear explants robustly and specifically induces hair cell marker expression in the GER, both *in vitro* ^39^ and *in vivo* ^1,40^. Second, Sox2, an early marker of inner ear prosensory regions, is robustly expressed in the GER ^41^ (**Figure 1A**). Third, unlike other neonatal cochlear floor cells, which are mitotically quiescent, the GER dynamically replaces cells until the middle of the second postnatal week in mice, as revealed by thymidine analog labeling ^42^. Fourth, gene expression profiling suggests that conditionally immortalized cochlear cells likely originated from GER cells ^43^. Fifth, when supporting cells surrounding inner hair cells were specifically ablated in neonatal mice, the neighboring GER cells replaced the lost cells ^7,12^. This replacement can also involve S-phase reentry and is accompanied by changes in gene expression following damage in the neighboring organ of Corti (**Figure 8**) ^8^. Finally, purified GER cells can give rise to proliferating otic progenitors that generate otic organoids (**Figure 1C**) ^9,44^. Together, these findings raise the possibility that regenerative mechanisms present in the neonatal GER may be reactivated in adult tissues to rekindle regenerative capacity.

To characterize the proliferative mechanisms of GER cells as a framework for future reprogramming of adult cochlear cells for hair cell regeneration, the first step of this study was to systematically define the gene expression profile of organoids at various stages of organoid formation. Our single-cell RNA-sequencing data obtained from GER-derived organoids revealed that *Lgals1* mRNA is highly enriched, while *Lgals3* and *Myc* mRNA are distinctly expressed during organoid formation at the onset of cell proliferation (**Figure 3C-E**).

Galectins are a family of proteins that bind to the β-galactoside-containing side chains of glycosylated proteins. They participate in diverse biological processes by mediating cell-cell and cell-matrix interactions and can initiate intracellular signaling ^19^. Galectins 1 and 3 have been implicated in cancer progression ^45,46^, which has motivated the recent application of the small-molecule inhibitors used in our screen (OTX008 and GB1107) as promising drug candidates for cancer and fibrosis treatments ^47^. Because galectins can mediate extracellular and cell-cell interactions, their growth-promoting effects need not require cell-autonomous co-expression with proliferation markers. In the inner ear, the role of galectin-1 is largely unexplored, aside from its reported expression in the inner and outer sulcus of the mature mouse cochlea ^2^. Myc, a multifunctional transcription factor and oncoprotein, along with Notch, has recently been shown to enable cell cycle entry in various mature cochlear floor cell types, including hair cells and supporting cells ^4^.

Galectin-1, galectin-3, and Myc inhibition dose-dependently suppressed organoid growth and formation (**Figures 4 and Figure 5**). The overexpression of galectin-1 and Myc, but not galectin-3, augmented organoid formation and cell growth (**Figure 6**). Together, the pharmacological and gain-of-function experiments support roles for galectin-1 and Myc in promoting organoid growth. The inhibitor experiments alone cannot distinguish effects on proliferation from cell survival, whereas the AAV studies link galectin-1 and Myc to cell-cycle activity.

Cochlear epithelial cells lose the ability to enter the cell cycle and form organoids following the onset of hearing in mice, around postnatal day 12 ^32^. Our finding that purified P14-17 cochlear epithelial cells can form organoids in response to Myc overexpression, but not galectin-1 (**Figure 7C-E**), demonstrates that selected genes from our organoid data can induce organoid-forming capacity in an EpCAM-enriched post-neonatal cochlear cell population. Prior studies showed that Myc can induce proliferation of adult cochlear supporting cells and hair cell regeneration *in vivo* ^4^. The identification of Myc in our GER-derived organoid single-cell RNA sequencing dataset, together with the increased cell proliferation in organoids grown from P14-P17 cochlear epithelium cells, provides proof of concept that GER-derived mechanisms can be leveraged to discover factors that promote cochlear hair cell regeneration in adult tissue. Because our organoid dataset is generated exclusively from GER cells, it provides a high-yield set of candidate genes that could drive a “GER-like” conversion of adult supporting cells for further exploration.

The *in vivo* damage response shared selected features with the pathways identified in the organoid system, although their spatial organization differed. Following damage, Lgals1 and Myc expression increased predominantly in the medial GER, whereas proliferating cells arose primarily in the lateral GER (**Figure 8 and Figure 9**). Galectin-1 protein expression broadened toward the lateral GER, suggesting that galectin-associated regulation of proliferation may not require cell-autonomous co-expression with proliferation markers. Inhibition of galectin-1 resulted in a reduction in the number of cycling cells within the GER: we propose that galectin-1-driven cell-cell and cell-matrix interactions, either directly or indirectly through systemic effects, are likely key initiators of GER proliferation, as predicted by our organoid model, where direct cell-cell contact plays a critical role in driving this process ^9^. Together, the alignment between our organoid and *in vivo* findings provides strong support for our strategy of utilizing organoid-based transcriptomic data to identify candidate genes required for cochlear epithelial cell regeneration *in vivo*.

An important translational consideration is that pathways capable of restoring proliferative competence overlap with pathways implicated in oncogenesis. Myc is a potent regulator of cell-cycle entry and a well-established proto-oncogene, whereas galectins 1 and 3 have been associated with tumor progression. Consequently, sustained or spatially unrestricted activation of these pathways would raise significant safety concerns. Our results do not establish galectins or Myc as immediately applicable therapeutic targets, but rather identify them as components of the molecular programs associated with cochlear progenitor growth. Future regenerative strategies would require tightly controlled and transient activation together with demonstration of appropriate cell-cycle exit and long-term safety.

In conclusion, this study identified galectin-1, galectin-3, and Myc as regulators of inner ear organoid growth by generating organoids from purified GER cells, integrating single-cell RNA sequencing, and functionally testing signaling pathways identified in proliferating organoid cells. Both galectin-1 and Myc effectively enhanced organoid growth. We then demonstrated the *in vivo* relevance of our organoid model and established it as a platform for identifying key effectors that enable the regeneration of cochlear epithelial cells.

## Materials and Methods

### Experimental Model and Study Participant Details

All animal care and procedures complied with the Guide for the Care and Use of Laboratory Animals published by the National Institutes of Health. Animal procedures were approved by Stanford University’s Institutional Animal Care and Use Committee and conducted per the NIH criteria described in ‘The Guide for the Care and Use of Laboratory Animals’.

### Cell Isolation

#### Neonatal cochlea

Temporal bones were removed from P2-4 mice, cochlear ducts were dissected, and the spiral ligament and stria vascularis were removed from each specimen. The tissues were rinsed twice with potassium- and magnesium-free PBS and incubated in a drop of 50 µL PBS mixed with 50 µL 0.25% Trypsin-EDTA (Gibco) at 37 °C and 5% CO_2_ for 10 min in a suspension culture dish (Greiner Bio-One). After adding 50 µL DMEM/F12 and 50 µL of a DMEM/F12-based mixture of soybean trypsin inhibitor (20 mg/mL) (Gibco) and DNAseI (2 mg/mL) (Stemcell Technologies), the cochlear duct tissue was mechanically dissociated with a pipette and passed through a 70 µm strainer to remove aggregates. The cells were subsequently passed through a 35 µm strainer for flow cytometric analysis and sorting.

#### P14-P17 cochlea

The cochleae were separated from the vestibular compartments, and the apex bone was removed from each specimen. Ten cochleae were collected into HBSS on ice, transferred to 700 μL thermolysin (0.5 mg/ml in DMEM), and gently vortexed at low speed, followed by incubation at 37 °C for 10 min. The sample was then centrifuged at 300 x g for 1 min, and the supernatant was removed. 700 μL Accutase was added, and the sample was gently mixed by vortexing at low speed, followed by incubation at 37 °C for 2 h. The sample was centrifuged at 300 x g for 1 min and the supernatant was removed. After washing once with PBS, 500 μL PBS was added, and the cells were mechanically dissociated by triturating with a low-retention 1000 μL pipette tip. Following centrifugation at 300 x g for 5 min and the removal of the supernatant, the cells were resuspended in MACS buffer (1:20 mixture of MACS BSA Stock Solution, Miltenyi Biotec / autoMACS Rinsing Solution, Miltenyi Biotec), and filtered with a 30 μm MACS SmartStrainer (Miltenyi Biotec) to remove bony fragments and aggregates. The cells were then subjected to magnetic cell separation.

### Magnetic Cell Separation

The dissociated and filtered cells were washed with 1 mL MACS buffer (1:20 mixture of MACS BSA Stock Solution, Miltenyi Biotec/ autoMACS Rinsing Solution, Miltenyi Biotec), and filtered with a 30μm MACS SmartStrainer (Miltenyi Biotec), followed by centrifuging at 300 x g for 10 min and the removal of supernatant. The cells were resuspended in 90 μL MACS buffer and incubated with 10 μL CD326 (EpCAM) MicroBeads (Miltenyi Biotec) at 4 °C for 15 min. The cells were washed with 2 mL MACS buffer, centrifuged at 300 x g for 10 min, and resuspended in 500 μL MACS buffer. The cells were applied to an MS Separation column (Miltenyi Biotec), pre-equilibrated with 3 mL MACS buffer, and placed into the MACS Separator (Miltenyi Biotec). The column was washed 3 times with 3 mL MACS buffer each. The column was taken out from the separator, and the enriched CD326-positive cells in the column were flushed with 3 mL MACS buffer into a 15 mL conical tube. After centrifugation at 300 x g for 3 min, the supernatant was removed. The cells were washed with 1 mL DMEM/F12 and centrifuged at 300 x g for 3 min. The cells were resuspended in the maintenance medium for culture (**Figure 7B and C**).

### Flow Cytometry

#### Sorting

Dissociated cochlear duct cells from Fgfr3-tdTomato/Sox2-GFP mice ^9,13^ were collected in a round-bottom tube and washed twice with phenol red-free DMEM/F12 medium (Gibco). For each wash, the cells were pelleted at 300 x g for 5 min and gently resuspended in medium. Sorting was performed on a BD ARIA II SORP flow cytometer using previously established procedures ^9,13^. Briefly, debris and doublets were excluded based on scatter and singlet parameters, respectively, and dead cells were eliminated using the CYTOX Red cell viability marker (Invitrogen). GFP-high and tdTomato-negative cochlear cells were sorted from Fgfr3-tdTomato/Sox2-GFP mice to collect GER cells for further culture (**Figure 1B**, **Figure 5**).

#### Analysis

Organoids were washed twice with potassium and magnesium-free PBS and incubated in a drop of 50 µL PBS mixed with 50 µL 0.25% Trypsin-EDTA (Gibco) at 37 °C and 5% CO_2_ for 10 min in a suspension culture dish (Greiner-bio-one). After adding 25 µL soybean trypsin inhibitor (40 mg/mL) (Gibco) and 75 µL DMEM/F12, the organoids were mechanically dissociated with a pipette. For the analysis described in **Figure S4C**, the cells were passed through a 35 µm strainer to remove aggregates and processed using flow cytometry with the parameters described above to quantitatively assess the GFP-positive cell population. For the analysis described in **Figure 6G**, dissociated organoid cells were incubated with Far Red Dead Cell Stain (Invitrogen) for 30 min at room temperature under light-protection, followed by fixation with 4% paraformaldehyde for 15 min. After each incubation step, cells were washed once with PBS and centrifuged at 300 x g for 5 min, and the supernatant was removed. Cells were then passed through a 35 µm strainer and subjected to debris and doublet discrimination by flow cytometry. The fluorescence of Far Red Dead Cell Stain was used to determine the proportion of dead and live cells.

### Organoid Culture

Cell seeding densities were kept consistent across experiments; Dissociated cochlear duct cells were seeded at 1.1 x 10^4^ cells per cm^2^ surface area (**Figure 4A, D, and G**; **Figure 6**, **Figure 7C**; **Figure S3**; **Figure S4B**). FACS-enriched GER cells were seeded at 2,000 cells per well in 96-well plates (**Figure 1C**; **Figure 5**).

To grow GER-organoids for single-cell RNA sequencing (**Figure 1B**), enriched GER cells were sorted into a 96-well suspension culture dish at 2000 cells/well filled with 200 µL/well maintenance medium. The maintenance medium consisted of DMEM/F-12 (Gibco) with 1x N-2 Supplement (Gibco), 1x B-27 Supplement (Gibco), and 100 µg/mL ampicillin.

Growth factors (20 ng/mL EGF, 10 ng/mL FGF2, and 50 ng/mL IGF1) and small molecules (3µM CHIR99021, 500 µM valproic acid, 100 µg/mL 2-phospho-L-ascorbic acid, and 2 µM TGFß receptor inhibitor 616452) were added ^5,13^. The cells were incubated for up to 7 days to allow organoid formation at 37°C and 5% CO_2_. Half of the medium was changed every other day. To grow organoids for evaluation of organoid growth (**Figure 4A, D, and G**, **Figures S3A-C**, **Figure S4B**, **Figure 6, and Figure 7B, C**) and immunohistochemistry (**Figure 3A and B**), dissociated cochlear duct cells were seeded in suspension culture dishes at 1.1 x 10^4^ cells per cm^2^ surface area in maintenance medium containing growth factors (20 ng/mL EGF, 10 ng/mL FGF2, and 50 ng/mL IGF1) and 3µM CHIR99021. The cells were cultured for 3 days (immunohistochemistry) or 7 days (evaluation of proliferation) at 37°C and 5% CO_2_. The same media composition was used for FACS-enriched GER cells (**Figure 5**). Here, the cells were seeded into a 96-well suspension culture dish at 2000 cells/ well filled with 200 µL medium and cultured for 7 days at 37°C and 5% CO_2_ to evaluate organoid growth. Organoid number and size were quantified using images captured under an inverted microscope by measuring organoids collected at the center of the field of view in individual wells of 48-well plates (**Figure 4B, E, and H**, **Figure 6E and I**, **Figure 7D**).

### Single-Cell Collection from GER-Derived Organoids (Figure 1C)

Organoids, harvested at culture days 1, 3, and 7, were individually collected with a small amount of medium with a low retention pipette into a well of a suspension culture dish (Greiner-bio-one) using an inverted microscope. The organoids were sequentially rinsed by transferring them into a 100 µL drop and then into a 50 µL drop of potassium and magnesium-free PBS. 50 µL of 0.25% Trypsin-EDTA (Gibco) was added, and the samples were incubated at 37 °C and 5% CO_2_ for 10-20 min. Subsequently, 25 µL of soybean trypsin inhibitor (40 mg/mL) (Gibco) and 75 µL of DMEM/F12 were added. The organoids were mechanically dissociated with a 200 µL low-retention pipette tip. The dissociated cells were transferred into a 1.5 mL low-adhesion tube and washed twice with DMEM/F12. Each wash was performed by centrifuging at 300 x g for 10 min and removing the supernatant. The dissociated cells were then transferred into a 500 µL drop of DMEM/F12 prepared in a cell-repellent culture dish (Greiner-bio-one, #628979) to prevent cell adhesion to plastic and minimize cell damage.

A micropipette puller (Sutter Instrument) was used to generate a 50-60 µm diameter glass micropipette from a glass capillary (World Precision Instruments). The micropipette was attached to the micropipette holder (Sutter Instrument) and connected with a clear plastic tube. Using an inverted microscope for visualization, single cells were aspirated into the glass capillary along with less than 0.5 µL of the medium. Each aspirated cell was individually released into a well of 96-well plates pre-filled with 4 µL lysis buffer (2U/μL RNase inhibitor, 0.2% (vol/vol) Triton X-100, 2.5μM modified Oligo-dT primer (5’-AAG CAG TGG TAT CAA CGC AGA GTA CTT TTT TTT TTT TTT TTT TTT TTT TTT TTT TVN - 3’). After collecting the cells, 96-well plates were immediately placed into a box filled with dried ice and then stored at -80°C.

### Single-Cell RNA-seq Library Preparation and Sequencing for GER-derived Organoid Cells

Library preparation and sequencing were conducted at the Stanford Functional Genomics Facility. Reverse transcription was performed with SMARTScribe Reverse Transcriptase (Clontech), followed by cDNA synthesis as described in the Smart-seq2 protocol ^15^. Synthesized cDNA was purified with AMPure XP beads (Beckman Coulter), quantified, and assessed for size distribution using the HS NGS Fragment Kit (1-6000bp) (Agilent) on a fragment analyzer (Advanced Analytical Technologies). The samples were concentration-normalized and tagmented using the Nextra XT (Illumina) sample preparation system, which adds cell-specific barcodes to the fragments. After PCR amplification and purification with AMPure XP beads, the cDNA library was quality-assessed using a Bioanalyzer 2100 (Agilent). Sequencing was performed on an Illumina NextSeq 500 using a high-output flow cell configuration with paired-end sequencing of 150 bp.

### Sequencing Data Processing for GER-derived Organoid Cells

The sequenced reads were aligned to the mouse reference genome release GRCm38/mm10 and the GENCODE annotation vM17 using custom scripts on the Sherlock Supercomputer Cluster (Stanford). The STAR aligner was used to map the raw sequencing reads, followed by RSEM for transcript quantification. Count matrices were loaded into R and processed with Seurat package version 5.0.1 ^16^. A total of 768 cells were sequenced from two batches of sequencing plates (384 cells x2 batches). Both datasets yielded 52,640 features each after removal of spike-in control RNAs (ERCCs). Features expressed in fewer than four cells were removed, leaving 18,952 and 17,426 genes, respectively, in these datasets. Cell-level quality control was performed by filtering out cells with fewer than 200 unique genes (nFeature_RNA >200) and cells with over 10% mitochondrial counts (percent.mt <10). The data was normalized with the scTransform function followed by integration in Seurat. The integrated dataset contained 658 cells and 18,587 features, which were clustered into 7 groups and visualized in a UMAP (**Figure 2A**). The CellTrails package (version 1.16.0) ^48^ was used to analyze the same datasets for quality control and normalization. The resulting dataset, comprising 646 cells and 23,473 features, yielded 13 clusters and was visualized using SWNE plots ^18^ (**Figure S1**). Transcription factors were identified using the AnimalTFDB 4.0 database ^49^ **(Table S1, S3, S4, and S5)**.

### Single-Cell Collection, Single-Cell RNA-seq Library Preparation and Sequencing, and Data Processing for Lgr5-DTR and Control Cochlear Cells (Figure 8A-H)

We used *Ki67^CreERT2/+^; Lgr5^DTR/+^; Rosa26R^tdTomato/+^* mice as the damage group and *Ki67^CreERT2/+^; Rosa26R^tdTomato/+^* littermates as controls, as described previously ^12^. *Ki67^CreERT2/+^; Rosa26R^tdTomato/+^* reporter expression was not used for cell enrichment or selection for sequencing. Cochlear cells were captured without reporter-based sorting, and epithelial populations were identified computationally after sequencing. Diphtheria toxin was administered intraperitoneally at P1 (4 ng/g), and tamoxifen was administered intraperitoneally at P3 (0.25 mg/g) to activate Ki67-CreERT2-mediated recombination. Cochleae were collected at P4 into ice-cold HBSS before incubation in 0.5 mg/mL Thermolysin (Sigma-Aldrich) at 37°C for 45 minutes. Tissues were then transferred to 300 µL Accutase and incubated for an additional 20 minutes at 37°C. After incubation, 300 µL of DMEM/F12 with 5% fetal bovine serum was added, and the tissue was dissociated into a single-cell suspension by 60x trituration at a 500 µL volume using a 1000 µL pipette tip. Remaining aggregates were removed by filtering the cells through a 40 µm mini strainer (pluriSelect) and cells were pelleted by centrifugation at 300 x g for 5 min at 4°C. The cell pellet was resuspended in 60 µL DMEM/F12 with 5% FBS and assessed for concentration and viability before capture using the 10x Genomics Chromium Single Cell 3’ Reagents v3.1 kit (10x Genomics) at the Stanford Functional Genomics Facility. Resulting single-cell libraries were sequenced on a NovaSeq 6000 (Illumina) using a 150bp paired-end read configuration.

Reads were aligned to the mouse reference genome mm39 with appended genome for the TdTomato-WPRE sequence using 10x Genomics Cell Ranger v6.0.0 with default parameters ^50^. Filtered digital gene expression matrices were used for downstream data analysis using Seurat v5.1.0 ^51^. Briefly, cell profiles were filtered based on the capture of sufficient unique counts and features, as well as a maximum of 10% mitochondrial gene content, and cell doublets were removed using DoubletFinder. Cell profiles from damage and control conditions were integrated using Harmony ^52^ and cells of interest were isolated based on the expression of known epithelial, IPhC and GER markers. Data are deposited in the Gene Expression Omnibus (GEO, accession number GSE291228).

### Immunohistochemistry

#### Organoid (Figure 3A and B)

Organoids were fixed with 4% paraformaldehyde for 30 min and washed three times for 10 min with PBS. The organoids were then permeabilized in 0.1% (w/v) TritonX-100 in PBS for 30 min, and blocked with 5% (v/v) heat-inactivated donkey serum, 1% (w/v) BSA, 0.1% (w/v) TritonX-100, and 0.02% (w/v) NaN_3_ in PBS for 1 h at room temperature. The organoids were then transferred from the plastic dish to a glass-bottom dish (Lab-Tek Chamber Slide, Thermo Fisher Scientific), which prevents permanent adhesion of the organoids to the dish’s bottom. The organoids were incubated with the primary antibodies in blocking buffer overnight at 4°C. The organoids were then washed three times for 15 min with PBS and incubated with secondary antibodies and DAPI at room temperature for 1 h in a buffer containing 0.1% (w/v) BSA, 0.1% (w/v) TritonX-100, and 0.02% (w/v) NaN_3_ in PBS. After washing three times for 10 min in PBS, the organoids were transferred to a glass slide with a spacer (Secure-Seal Spacer, Invitrogen), sealed with ProLong Diamond Antifade mounting medium (Invitrogen), and a coverslip.

#### P14-P17 cochlea (Figure 7A)

The inner ear was dissected from P14-17 FVB/NJ mice and fixed with 4% paraformaldehyde for 2 h. After three 15 min washes with PBS, the tissue was incubated in 0.5% EDTA, pH 8.0 overnight at 4°C with gentle agitation for decalcification. Tissues were then embedded in 4% Low Melt Agarose (Bio-Rad) at 65°C in dH_2_O. The tissue blocks were cooled until the agarose solidified, and 100 μm sections were cut using a vibratome (VT100S, Leica). Permeabilization, blocking, immunostaining, and mounting were performed as described above.

#### Whole cochlear duct (Figures 8I, Figure 9, Figure S5, Figure S6)

The temporal bones were dissected from P4 mice and fixed with 4% paraformaldehyde overnight at 4°C with gentle agitation. After washing with PBS three times for 15 min, cochlear ducts were dissected and attached to 10 mm coverslips with Cell-Tak (Corning). Permeabilization, blocking, and immunostaining were performed as described above. Each tissue-mounted coverslip was placed onto a glass slide, and sealed with ProLong Diamond Antifade mounting medium (Invitrogen) and another coverslip.

Primary antibodies used: goat anti-galectin-1 (1:50, R&D Systems, Cat# AF1245), rabbit anti-galectin-3 (1:100, Proteintech, Cat# 14979-1-AP), rat anti-Ki-67 (1:250, Thermo Fisher Scientific, Cat# 14-5698-80), rabbit anti-Myosin-Vlla (1:1000, Proteus, Cat# 25-6790), Rat APC anti-mouse CD326 (Ep-CAM) (1:50, BioLegend, Cat# 118214), rabbit anti-fatty acid binding protein 7 (FABP7) (1: 200, Abcam, ab32423). Secondary antibodies used: Donkey anti-rat Alexa Fluor 488 (1:250, Thermo Fisher Scientific, Cat# A-21208), Donkey anti-goat Alexa Fluor 647 (1:250, Thermo Fisher Scientific, Cat# A-21447), Donkey anti-rabbit Alexa Fluor 594 (1:250, Thermo Fisher Scientific, Cat#A-21207).

### Confocal Microscopy

Cochlear sections and organoids were imaged with an LSM880 confocal laser scanning microscope (ZEISS) and processed with ZEN (ZEISS) and Fiji software.

### Viable Cell Mass Assay

The CellTiter 96 Aqueous One Solution Cell Proliferation Assay System (Promega) was used to compare cell mass increases in organoid cultures. 20 μL of CellTiter 96 Aqueous One Solution reagent was added per 100 μL of culture medium and incubated at 37°C and 5% CO_2_ for 4 h. Absorbance at 490 nm was measured using a plate reader (Infinite M1000, Tecan). At least three independent experiments were conducted (**Figure 4C, F, and I**, **Figure 5B**, **Figure 6F and J**, **Figure 7E**).

### qPCR

The medium was removed from the organoids grown in 12-well suspension culture dishes, and the organoids were dissolved in 100 μL TRIzol Reagent (Invitrogen) per well. An additional 100 μL of 100% ethanol was added to each well, then mixed with the sample solution. The samples were loaded onto Direct-zol RNA Miniprep (Zumo Research) columns, and RNA extraction was performed according to the manufacturer’s protocol. The extracted RNA was reverse-transcribed using the SuperScript IV First-Strand Synthesis System (Invitrogen). All cDNA synthesized from 10 ng RNA was used in a single qPCR amplification session using Maxima SYBR Green/ROX qPCR Master Mix (2X) (Thermo Scientific) and the CFX96 Real-Time System (Bio-Rad) (**Figure 6B and C**).

### AAV-Vector Generation and Virus Production

The plasmid pAAV-CAG-EBFP2-P2A-EGFP (Addgene) was used as the backbone of AAV vector constructs. The control EGFP- and lgals1-encoding plasmids were produced using restriction digestion and ligation by replacing the EBFP2-P2A-EGFP sequence individually with PCR-amplified EGFP or lgals1 sequences. Similarly, lgals3- and myc-encoding plasmids were produced by replacing the EBFP sequence with lgals3 or myc. EGFP was further replaced with the mCherry sequence in the myc-encoding plasmid (**Figure S4A**). *Lgals1*, *lgals3*, and *myc* cDNA were generated from total RNA, extracted with the Direct-zol RNA Miniprep kit (Zumo Research) from organoids grown from whole cochlear duct cells of FVB/NJ mice. Reverse transcription utilized the SuperScript IV First-Strand Synthesis System (Invitrogen), and was followed by PCR with specific primers for lgals1, lgals3, and myc. The PCR products were subjected to electrophoresis, followed by DNA purification, restriction digestion, and subcloning. The resulting plasmids were sequenced to confirm correct cDNA inserts (GENEWIZ). Plasmids were purified from XL10-Gold *E. coli* (Agilent Technologies) with EndoFree Plasmid Maxi Kits (Qiagen) for viral production. Viral production involved transfecting HEK293 cells with the individual plasmids, a helper plasmid, and the Rep/Cap plasmid encoding serotype DJ protein. Viral production and purification were conducted by Stanford University’s Gene Vector and Virus Core facility.

### *In vivo* mouse models

For single-cell RNA sequencing (**Figure 8A-H**), *Ki67^CreERT2/+^; Lgr5^DTR/+^; Rosa26R^tdTomato/+^*(damage) and *Ki67^CreERT2/+^; Rosa26R^tdTomato/+^* (control) mice were used. Diphtheria toxin (DT) dissolved in saline (4 ng/g) was administered intraperitoneally (IP) at P1. For Cre recombination, tamoxifen dissolved in corn oil (0.25 mg/g) was administered at P3, the time point of peak proliferation post-damage ^12^. Cochlear sensory epithelia were dissected at P4. For OTX008 treatment (**Figures 8I**, **Figure 9**, **Figure S5**, **Figure S6**), *Lgr5^DTR/+^* (damage) and control mice were used. After DT administration at P1, OTX008 (MedChemExpress) dissolved in 10% ethanol and 90% saline was administered intraperitoneally at P2 and P3 (10 mg /kg each time), or only at P2 (10 mg /kg), and cochlear sensory epithelia were dissected at P4.

### Quantification and Statistical Analysis

Statistical details for individual experiments are provided in the figure legends. Statistical analyses were performed in R version 4.2.0 and GraphPad Prism version 10.2.2.

## Data availability

All sequencing datasets generated in this study have been deposited in the National Center for Biotechnology Information (NCBI) Gene Expression Omnibus (GEO) under the accession numbers GSE326100 and GSE291228. The data are also accessible at gEAR, a gene Expression Analysis Resource (https://umgear.org/) ^53^. Further information and requests for reagents may be directed to the Lead Contact, Stefan Heller.

## Code availability

Single-cell RNA sequencing data were analyzed using R (v4.5.0) with publicly available R packages, following existing vignettes and code. No custom code was generated in this study. Details of the R packages and software used are provided in the Materials and Methods.

---

**Table S1:** Marker genes for clusters 1-7 in GER-derived organoids.

**Table S2:** Ingenuity pathway analysis results – top 10 pathways.

**Table S3:** Differentially expressed genes at the onset of proliferation.

**Table S4:** Marker genes for CellTrails clusters in GER-derived organoids.

**Table S5:** Differentially expressed genes at the onset of proliferation for CellTrails analysis.

**Table S6:** Ingenuity pathway analysis results for CellTrails analysis – top 10 pathways.

**Table S7:** Differentially expressed genes in organ of Corti cells.

**Table S8:** Differentially expressed genes in *in vivo* damage model (Lgr5-DTR) and control.

## Acknowledgments

We thank S. Sim and M.C. Yee for their invaluable support with single-cell sequencing, and the current and previous members of the Heller laboratory for their helpful discussions and manuscript review. We are grateful to the FACS Core of the Institute for Stem Cell Biology and Regenerative Medicine and the Gene Vector and Virus Core of Stanford University. We thank Stanford Functional Genomics Facility, which is supported by NIH grants S10OD018220 and 1S10OD021763, as well as the OHNS microscopy core facility, for their assistance with imaging. M.K. was supported by a grant from the National Institutes of Health R21DC020271, the Japan Society for the Promotion of Science, the Uehara Memorial Foundation, and the Soda Toyoji Memorial Foundation. We also acknowledge the generous support of the Hearing Restoration Project Consortium of the Hearing Health Foundation, the Stanford Initiative to Cure Hearing Loss, and the National Institutes of Health grants R21DC019910 (S.H), R01DC021110 and R01DC016919 (A.G.C.), K08DC019683 and R21DC022058 (T.A.J).

## Author contributions

Conceptualization, M.K. and S.H.; methodology, M.K., T.A.J; formal analysis, M.K.; investigation, M.K., P.K.L., S.BB., T.A.J, M.S.M., J.M.A., A.G.C., S.H.; visualization, M.K., M.S.M.; funding acquisition, M.K., T.A.J., S.H., A.G.C.; project administration, M.K., S.H.; supervision, M.K., S.H.; writing – original draft, M.K., P.K.L, S.H.; writing – review & editing, M.K., S.H., T.A.J., A.G.C., J.M.A.

