## Supplemental Figures for "Galectin and Myc enable cochlear progenitor expansion in vitro and in vivo"

Marie Kubota *et al.*

**This PDF file includes:**

Figs. S1 to S6

**Other Supplementary Materials for this manuscript include the following:**

Table S1 to S8

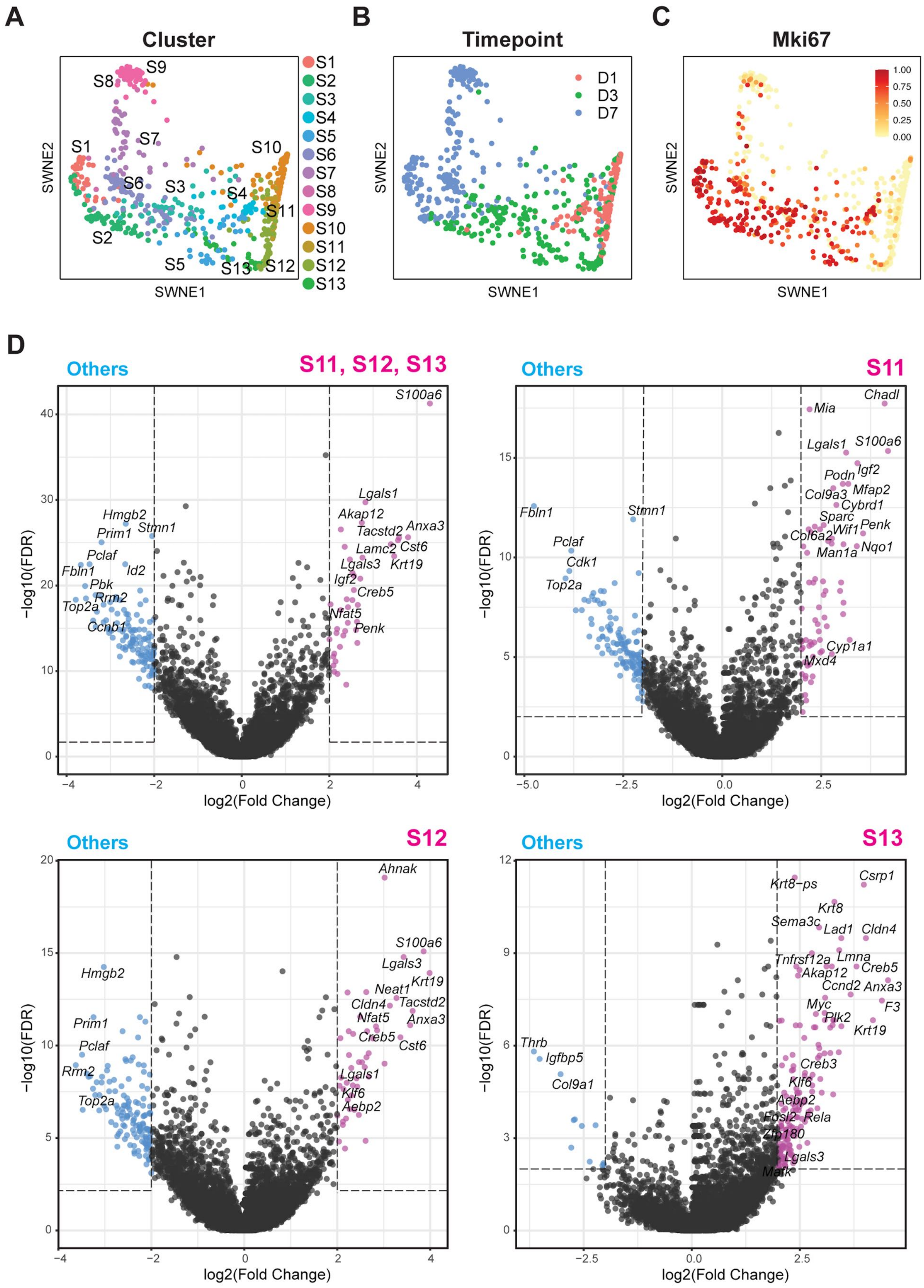

A
Cluster
S1
S2
S3
S4
S5
S6
S7
S8
S9
S10
S11
S12
S13
SWNE1
SWNE2
B
Timepoint
D1
D3
D7
C
Mki67
1.00
0.75
0.50
0.25
0.00
D
Others
S11, S12, S13
S11
S12
S13
log2(Fold Change)
−log10(FDR)
S100a6
Lgals1
Akap12
Tacstd2
Anxa3
Lamc2
Cst6
Lgals3
Krt19
Igf2
Creb5
Nfat5
Penk
Hmgb2
Prim1
Stmn1
Pclaf
Fbln1
Id2
Pbk
Rrm2
Top2a
Ccnb1
Chadl
Mia
Podn
Col9a3
Mfap2
Cybrd1
Sparc
Wif1
Col6a2
Man1a
Nqo1
Cdk1
Cyp1a1
Mxd4
Ahnak
Neat1
Cldn4
Klf6
Aebp2
Csrp1
Krt8−ps
Krt8
Sema3c
Lad1
Tnfrsf12a
Lmna
Ccnd2
Myc
Plk2
F3
Creb3
Fosl2
Rela
Zfp180
Matk
Thrb
Igfbp5
Col9a1

**Fig. S1. Single-cell RNA sequencing data analysis of GER cell-derived organoids using CellTrails.**

(A) Single-cell RNA sequencing data of GER-organoid cells were analyzed with CellTrails. Thirteen distinct clusters were identified and visualized with Similarity Weighted Nonnegative Embedding (SWNE). SWNE features encompass potential trajectory information, enabling us to track time points along GER-organoid development. (B) Timepoint visualization of the organoid cells harvested on days one, three, and seven (D1, D3, D7). (C) Expression of mRNA for the proliferation marker Kiel 67 (*Mki67*) was associated with cells in clusters S1-S6. (D) Volcano plot showing differentially expressed genes between the cells at the onset of proliferation (clusters S11, 12, and 13) and the remaining cells (clusters S1-S10) (top, left). The other three volcano plots show the differentially expressed genes in individual clusters S11 (top, right), S12 (bottom, left), and S13 (bottom, right). Thresholds are indicated with dotted lines: thresholds are set to log2-fold change > |2| and -log10(FDR) > 2.  Wilcoxon Rank Sum test was used for the comparison.

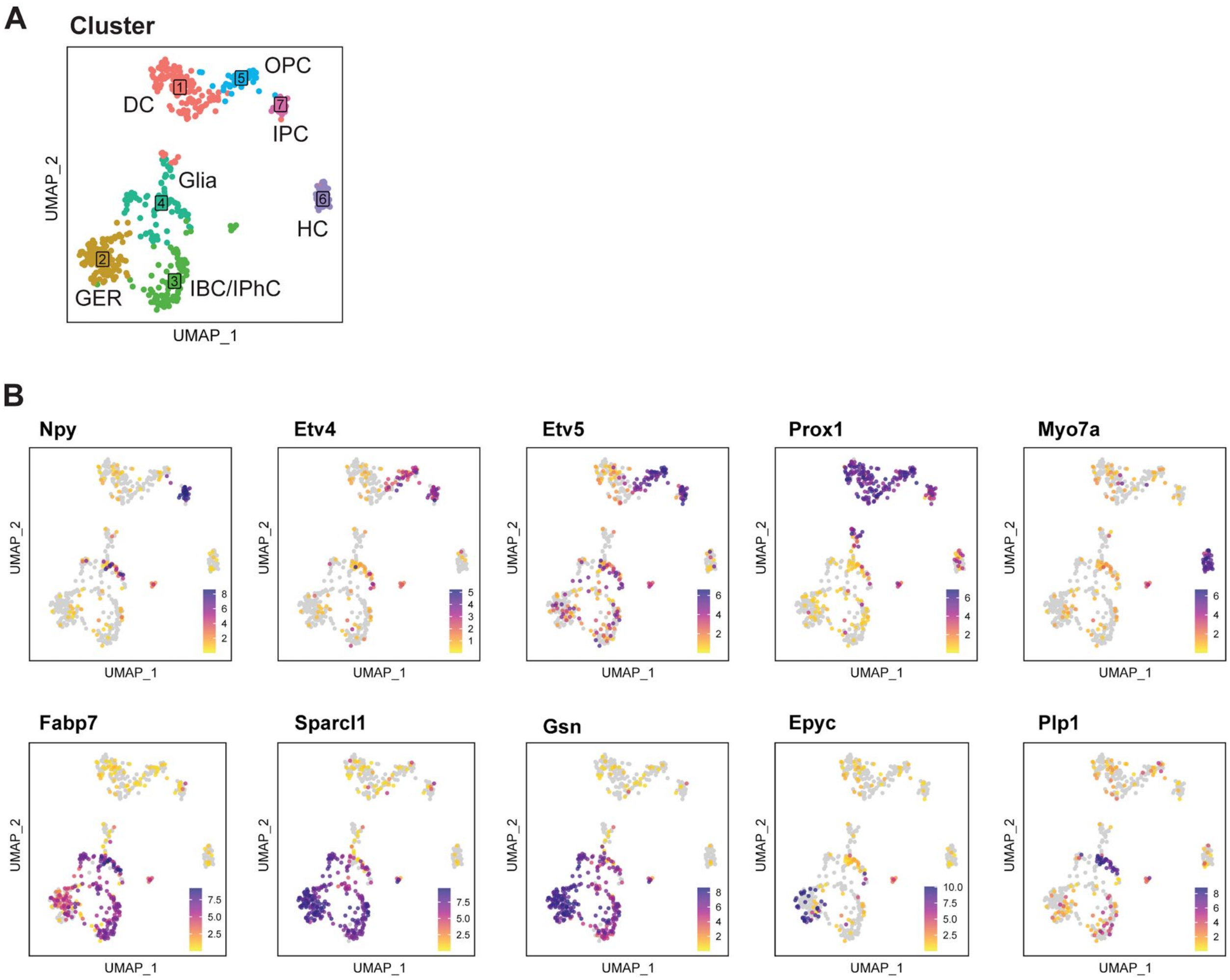


**Fig. S2. Cell type identification in single-cell data from cochlear floor cells.**

(A) Single-cell RNA-sequencing data obtained from P2 cochlear floor cells were subjected to cluster analysis using the Seurat package. Seven distinct clusters were identified. Shown are 548 FACS-isolated cells from Fgfr3-Cre-ERT2/Ai14-tdTomato/Sox2-GFP transgenic mice (ref). The individual clusters are labeled by the cell subtype. DC = Deiters' cells, GER = greater epithelial ridge, IBC = inner border cells, IPH = inner phalangeal cells, OPC = outer pillar cells, IPC = inner pillar cells, HC = hair cells. (B) Expression of known marker genes, projected into the UMAP plot: Npy = Inner pillar cells, Etv4 and Etv5 = pillar cells, Prox1 = pillar cells and Deiters' cells, Myo7a = hair cells, Fabp7 = inner phalangeal cells, Sparcl1 = GER, Gsn = GER, Epyc = medial GER, and Plp1 = glia.

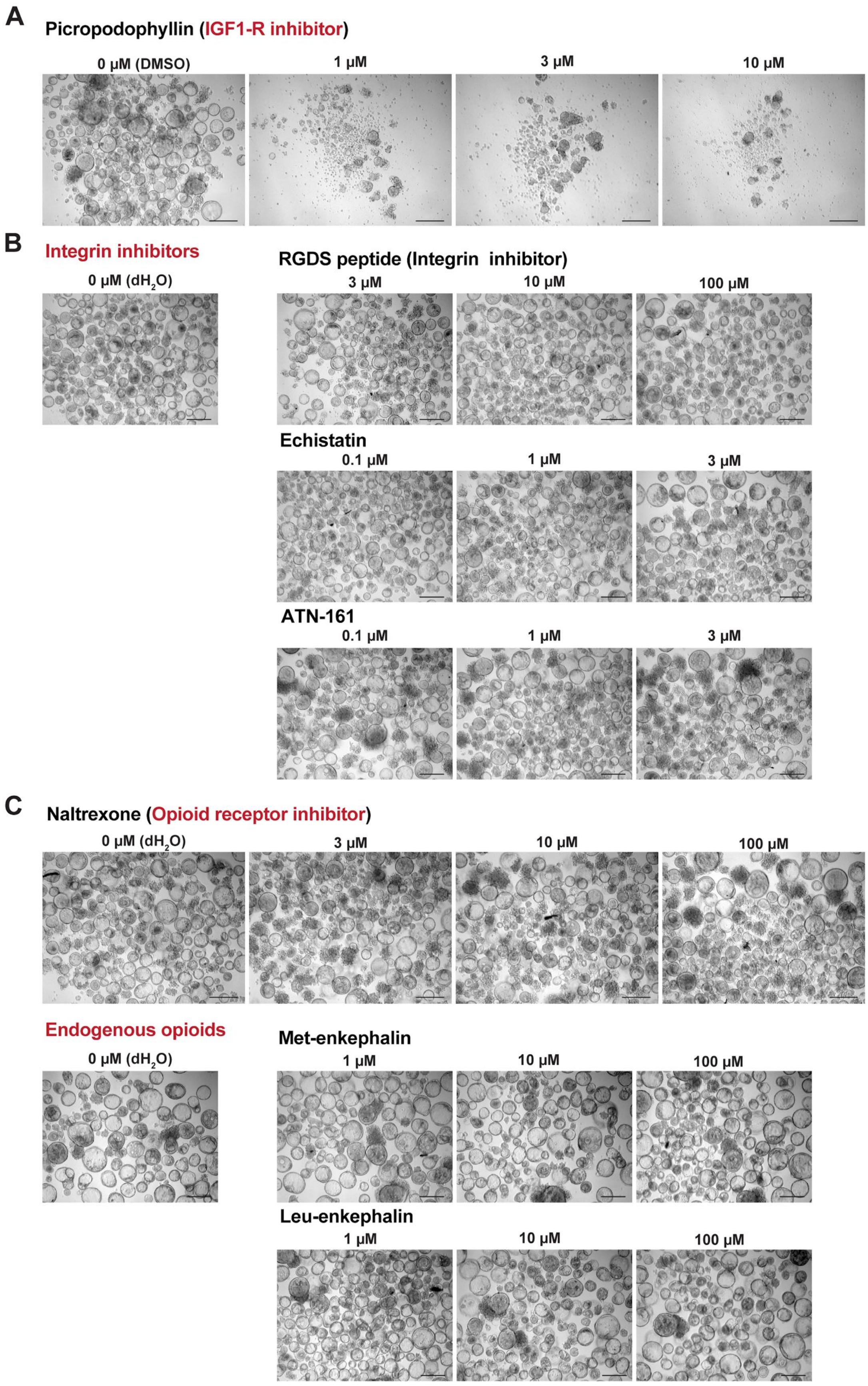
A
Picropodophyllin (IGF1-R inhibitor)
0 µM (DMSO)
1 µM
3 µM
10 µM
B
Integrin inhibitors
0 µM (dH2O)
RGDS peptide (Integrin inhibitor)
3 µM
10 µM
100 µM
Echistatin
0.1 µM
1 µM
3 µM
ATN-161
0.1 µM
1 µM
3 µM
C
Naltrexone (Opioid receptor inhibitor)
0 µM (dH2O)
3 µM
10 µM
100 µM
Endogenous opioids
0 µM (dH2O)
Met-enkephalin
1 µM
10 µM
100 µM
Leu-enkephalin
1 µM
10 µM
100 µM

**Fig. S3. Organoid formation assay screen with additional modulators.**

Organoids generated from whole cochlear duct cells of P2 FVB/NJ mice after seven days in culture. (A) Dose-dependent inhibition of organoid formation with picropodophyllin (IGF1-R inhibitor). Scale bar: 200 µm. (B) RGDS peptide, echistatin, and ATN-161 (integrin inhibitors) do not inhibit organoid formation. Scale bar: 200 µm. (C) Naltrexone (opioid receptor inhibitor), met-enkephalin, and leu-enkephalin (endogenous opioids) neither inhibit nor enhance organoid formation. Scale bar: 200 µm.

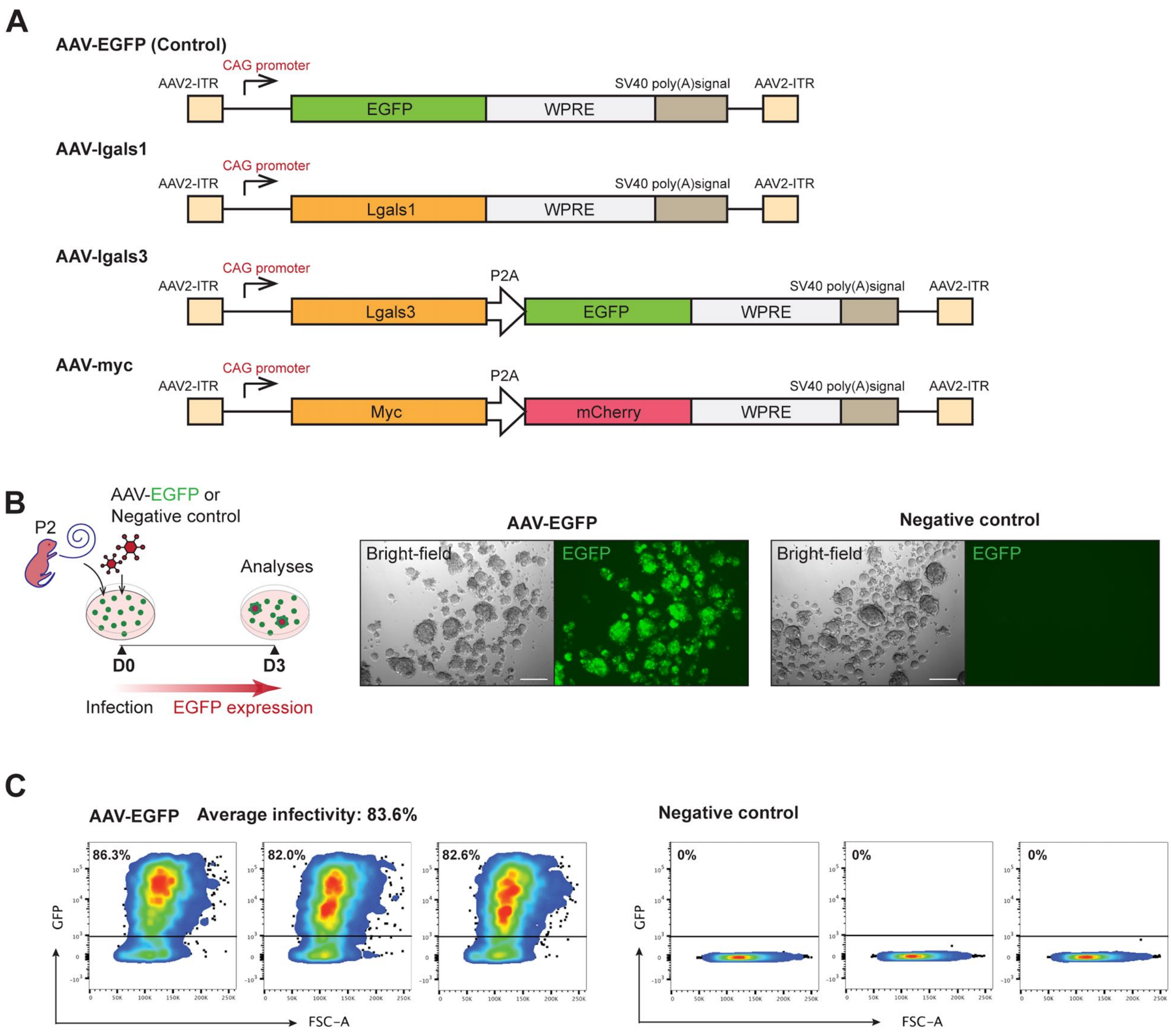


**Figure S4. AAV-mediated overexpression of genes in organoids.**

(**A**) AAV vector maps are shown. EGFP and Lgals1 coding sequences were cloned individually into the plasmid backbones. Lgals3 and Myc coding sequences were fused to EGFP and mCherry, respectively, with a P2A sequence separating them, and then cloned into the proviral plasmid backbone. (**B**) (Left) Schematic illustration of the organoid formation assay timeline using dissociated P2 FVB/NJ mouse cochlear duct cells infected with AAV-EGFP or control (medium only). (Right) Fluorescent microscopic images showing bright-field and green channel (EGFP) images of day 3 organoids infected with AAV-EGFP or control. Scale bars: 200 µm. (**C**) FACS analysis of dissociated organoid cells after 3 days in culture. On average, 83.6% of the cells expressed EGFP in the organoid infected with AAV-EGFP (N = 3).

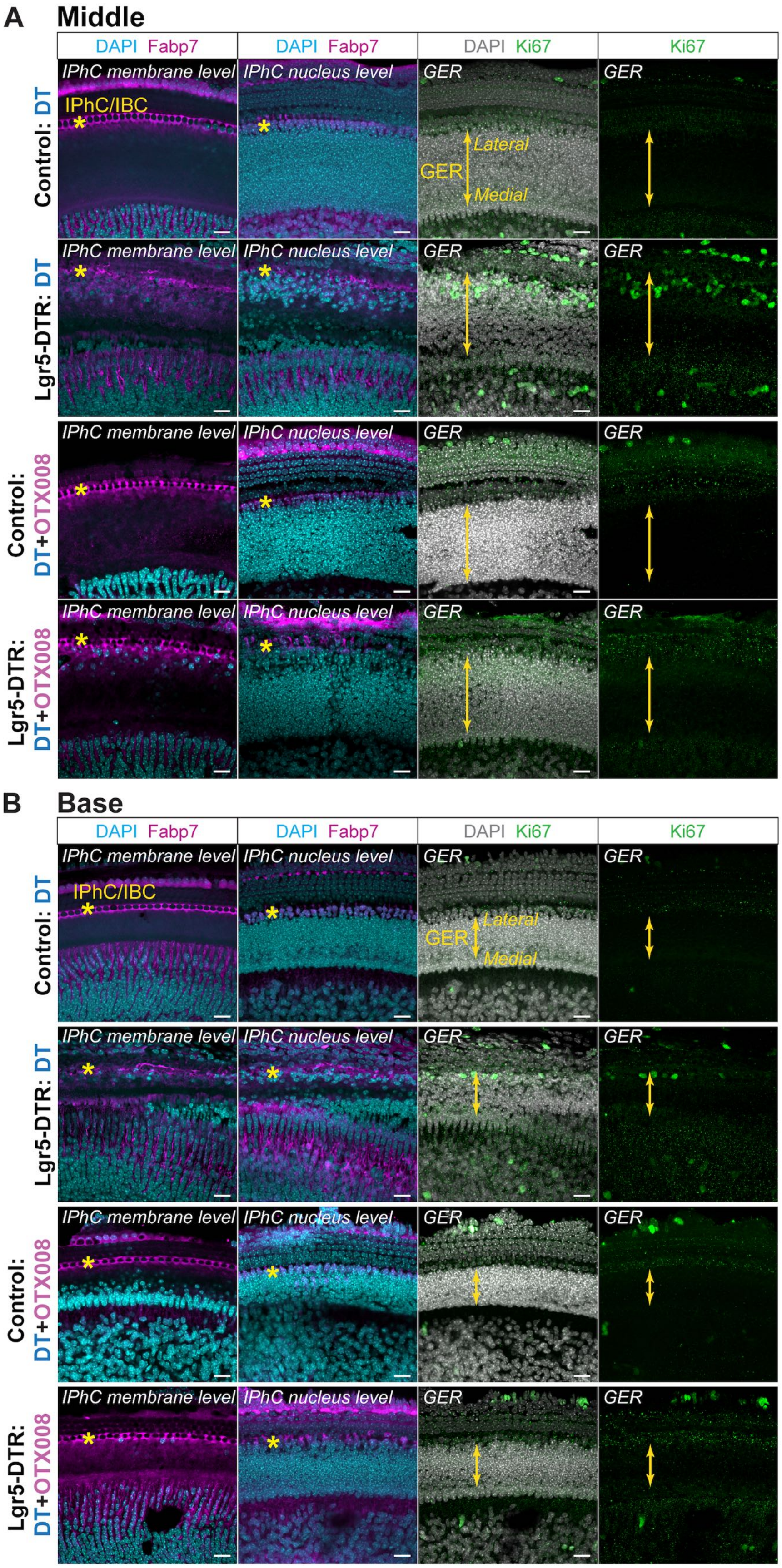
A
Middle
DAPI Fabp7
DAPI Fabp7
DAPI Ki67
Ki67
Control: DT
Lgr5-DTR: DT
Control: DT+OTX008
Lgr5-DTR: DT+OTX008
IPhC membrane level
IPhC nucleus level
GER
IPhC/IBC
*
Lateral
Medial
B
Base
DAPI Fabp7
DAPI Fabp7
DAPI Ki67
Ki67
Control: DT
Lgr5-DTR: DT
Control: DT+OTX008
Lgr5-DTR: DT+OTX008
IPhC membrane level
IPhC nucleus level
GER
IPhC/IBC
Lateral
Medial

**Figure S5. *In vivo* administration of galectin-1 inhibitor OTX008 at P2 and P3.**

Immunohistology of P4 cochleae obtained from control and Lgr5-DTR (*Lgr5*$^{DTR/+}$) mice treated with DT at P1, representing intact and damage controls, respectively, and from control and Lgr5-DTR mice treated with OTX008 (10 mg/kg per injection) at P2 and P3 after DT treatment at P1 (DT+OTX008). Stacked images of the middle (**A**) and basal (**B**) turns of the cochleae are presented at three levels: IPhC membrane, IPhC nuclei, and GER (scale bars: 20 μm). Asterisks indicate IPhCs and IBCs. Solid yellow arrows indicate the GER.

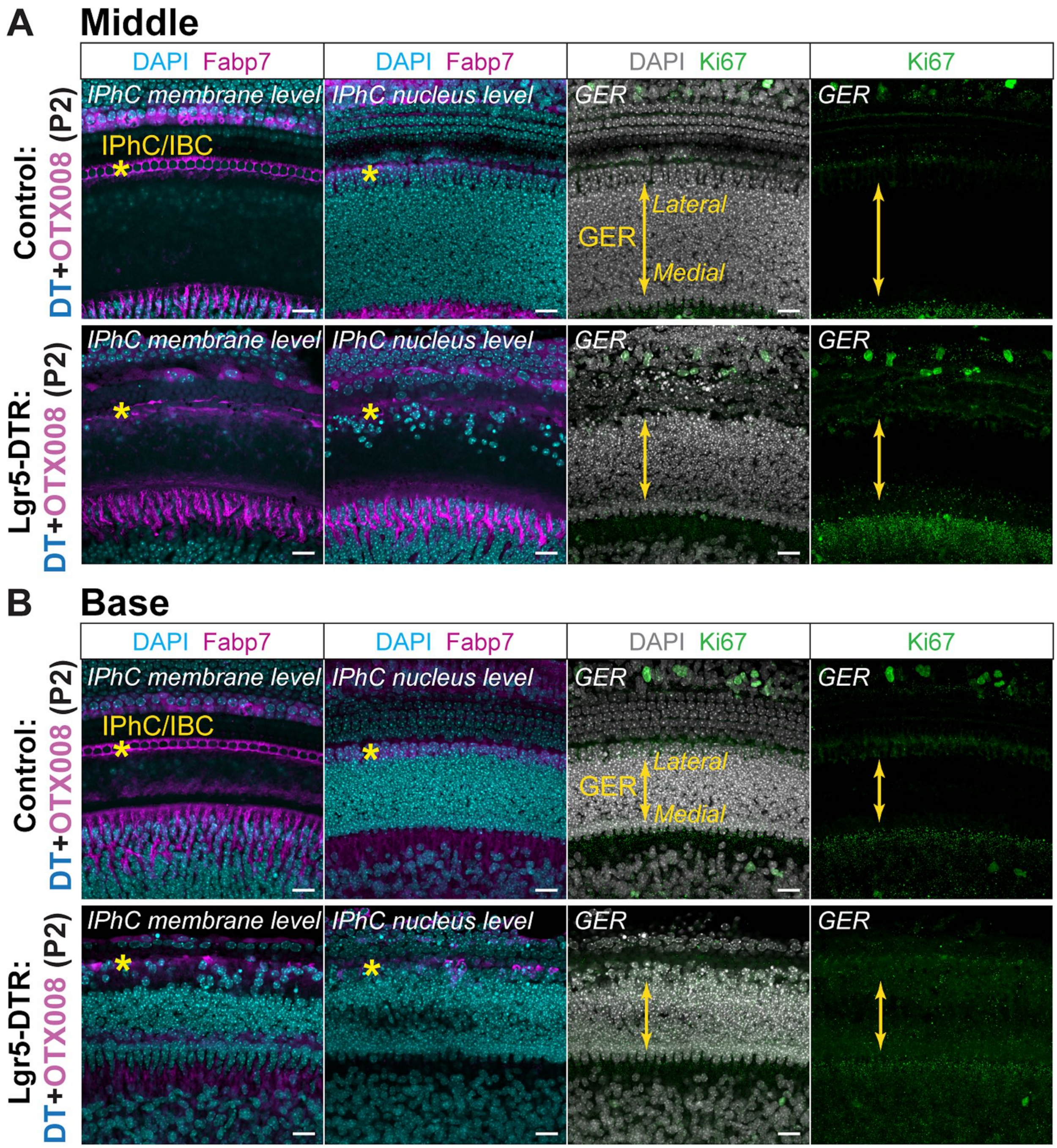


**Figure S6. *In vivo* administration of galectin-1 inhibitor OTX008 at P2.**

Immunohistology of P4 cochleae obtained from control and Lgr5-DTR mice treated with OTX008 (10 mg/kg per injection) only at P2 after DT at P1 (DT+OTX008 (P2)). Stacked images of the middle (**A**) and basal (**B**) turns of the cochleae are presented at three levels: IPhC membrane, IPhC nuclei, and GER (scale bars: 20 μm). Asterisks indicate IPhCs and IBCs. Solid yellow arrows indicate the GER.
